# The hearing-essential intracellular domain of PCDH15 reveals a new layer of auditory mechanotransduction

**DOI:** 10.64898/2026.09.17.752191

**Authors:** Jana Van de Velde, Ruxandra Bighiu, Nabanita Mandal, Karine Lapouge, Marie Skepö, Yann G.-J. Sterckx, Christine Petit, Vincent Van Rompaey, Pieter Van Wielendaele

## Abstract

Current models of mechanotransduction in inner-ear hair cells explain how force is transmitted through the extracellular tip link but provide little insight into how force is propagated beneath the plasma membrane. Although the CD2 isoform of PCDH15 has been shown to be essential for hearing in mature mammalian hair cells, its structural and mechanical properties have remained largely unknown. Here, we combine computational sequence analysis, orthogonal biophysical characterization and small-angle X-ray scattering (SAXS) to show that the hearing-essential CD2 intracellular domain is an intrinsically disordered region (IDR) that behaves as a highly expanded acidic polyampholyte. CD2 remains predominantly monomeric in solution and occupies a larger conformational space at physiological-like ionic strength, whereas inclusion of the isoform-shared common region increases self-association. CD2 also contains a conserved regulatory interaction platform, providing a potential link between its polymer properties and cellular regulation. Together, these findings identify the CD2 intracellular domain as a polymer with biophysical properties consistent with a mechanically responsive structural element. Furthermore, they provide a framework for investigating how intracellular polymer mechanics may contribute to force transmission and adaptation in auditory mechanotransduction.

## 2 Introduction

Sound is converted into electrical signals by mechanotransduction (MET) channels located at the tips of sensory hair-cell stereocilia^1^. Mechanical force is transmitted to these channels through the tip link, a heterotypic filament formed by cadherin-23 (CDH23) at its upper end and protocadherin-15 (PCDH15) at its lower end (Fig. 1a)^2^. The extracellular domain of PCDH15 has been extensively studied as a candidate gating spring, the compliant element that transmits force to the MET channel while allowing channel opening in response to mechanical stimulation^2–4^. Consequently, many mechanical models of auditory mechanotransduction have focused on the extracellular tip-link complex. More recent models have also emphasised force transmission through the intracellular MET-channel complex centred on TMC1, TMIE and CIB2^5–8^. Despite these advances, how the large intracellular region of PCDH15 contributes to the mechanical and regulatory behaviour of the mechanotransduction complex remains poorly understood.

**Figure 1.**
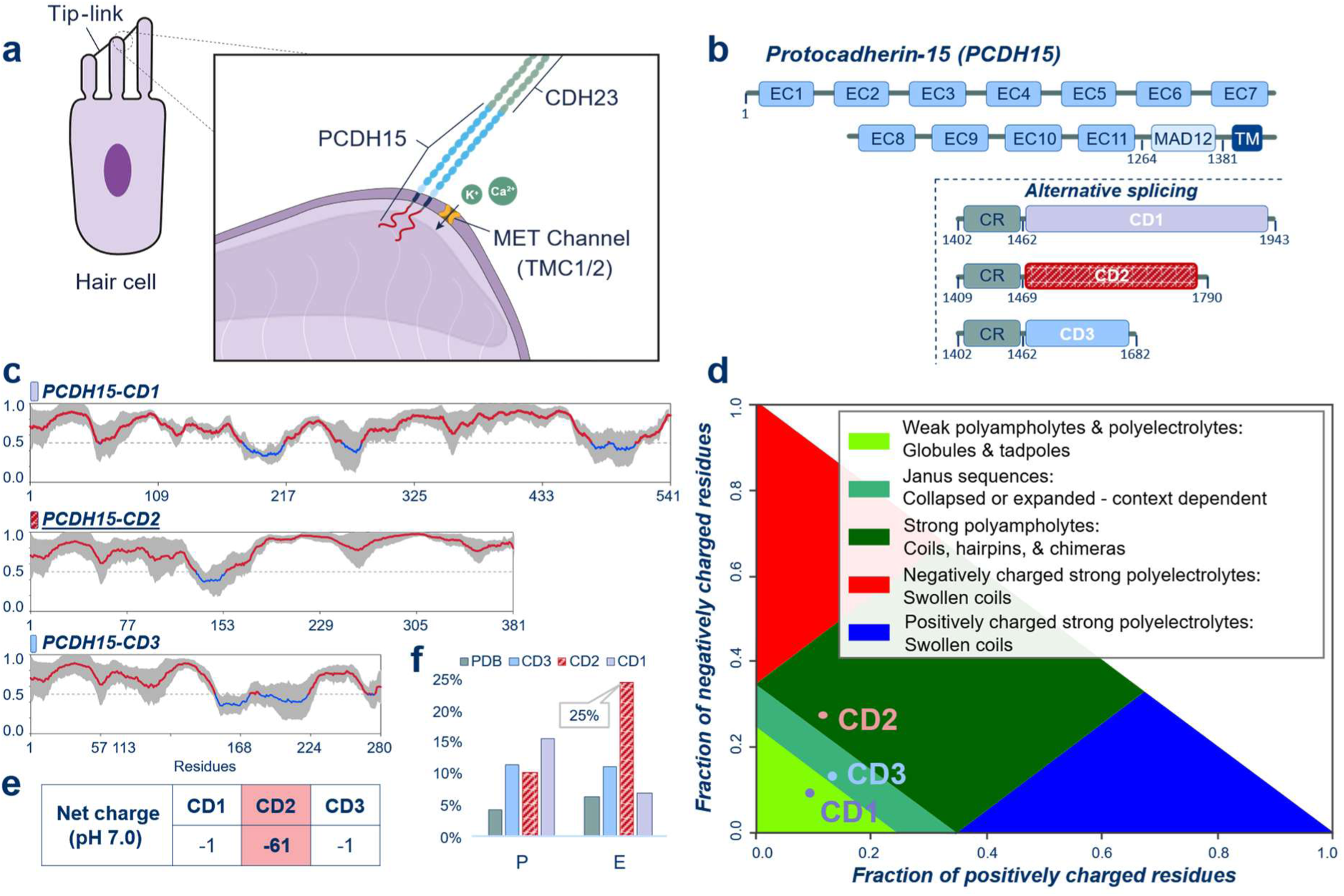
Sequence analysis identifies distinct polymer properties among the PCDH15 intracellular isoforms. **(a)** Schematic representation of a sensory hair cell and the lower tip-link insertion, showing PCDH15, cadherin-23 (CDH23) and the mechanotransduction (MET) channel. **(b)** Domain organization of protocadherin-15 (PCDH15), showing the extracellular cadherin (EC) repeats, membrane-adjacent domain (MAD12), transmembrane (TM) domain, and cytoplasmic common region (CR), followed by the alternatively spliced CD1-, CD2- or CD3-specific regions. **(c)** Consensus intrinsic-disorder predictions for the three PCDH15 cytoplasmic isoforms generated using RIDAO. Disorder scores range from 0 to 1, with scores above 0.5 indicating a predominant disorder propensity. **(d)** Das–Pappu phase diagram generated with CIDER showing the predicted polymer regimes of the three intracellular isoforms. PCDH15-CD1, PCDH15-CD2 and PCDH15-CD3 occupy distinct regions of the diagram corresponding to the globule/tadpole, strong polyampholyte and Janus sequence regimes, respectively. **(e)** Predicted net charge at pH 7.0 for the three PCDH15 cytoplasmic isoforms. **(f)** Relative abundance of proline (P) and glutamate (E) in the three PCDH15 cytoplasmic isoforms.

PCDH15 is a single-pass transmembrane protein comprising 11 extracellular cadherin (EC) repeats followed by a membrane-adjacent domain (MAD12), a transmembrane (TM) helix and a C-terminal cytoplasmic domain (CD) extending several hundred amino acids into the cytoplasm (Fig. 1b)^3,9^. PCDH15 forms a parallel cis-homodimer, an architecture that is important for mechanotransduction^10^. The protein is expressed as three alternatively spliced isoforms, each containing different CDs: PCDH15-CD1, PCDH15-CD2 and PCDH15-CD3, which share a common region but differ in their C-terminal sequences ^11–13^. Remarkably, loss of PCDH15-CD2 causes profound hearing loss in mouse models, whereas loss of CD1 or CD3 does not ^11,12^. This selective requirement suggests that the CD2 intracellular domain performs a unique function that cannot be replaced by the other splice variants. The structural and biophysical properties of the intracellular region of PCDH15 remain almost completely uncharacterised. Its structural organisation has not been established, making it unclear whether this domain functions simply as a molecular scaffold or whether its physical properties contribute directly to force transmission during mechanotransduction. Inspection of publicly available prediction resources suggests that the cytoplasmic regions of all three PCDH15 isoforms are intrinsically disordered. Intrinsically disordered proteins and regions (IDPs and IDRs, respectively) do not adopt a well-defined three-dimensional structure but are instead best described by a conformational ensemble containing various highly dynamic conformers that rapidly interconvert from one form to another. The mechanical properties of an IDP/IDR and the conformational space occupied by its conformers are determined largely by amino acid composition, charge distribution and the surrounding solution^14,15^. For the different PCDH15 CD isoforms, the sequence-based *in silico* predictions have not been experimentally validated, and they do not reveal how the intracellular domain behaves in solution or whether it possesses the physical properties required for a mechanical role. Although IDPs are increasingly recognised as important structural and regulatory elements in many biological systems, this perspective has rarely been applied to auditory mechanotransduction. While IDRs have been implicated in other auditory proteins, including prestin and harmonin, the structural and polymer-physical properties of the tip-link machinery itself remain largely unexplored^16,17^.

Here, we combine computational sequence analysis with orthogonal biophysical characterisation and small-angle X-ray scattering (SAXS) to determine the structural and physical properties of the intracellular domain of PCDH15 CDs. Computational analyses indicate that all three CD isoforms are intrinsically disordered, but that only the hearing-essential CD2 isoform possesses the sequence composition of a strongly acidic polyampholyte. Consistent with this prediction, recombinant CD2 adopts an exceptionally expanded conformation that increases further at physiological-like ionic strength and contains a highly conserved regulatory interaction platform centred on a candidate calcineurin docking site, an extensive phosphorylation cluster and a C-terminal PDZ-binding motif. Together, these findings identify the hearing-essential CD2 intracellular domain as an unusually expanded, salt-responsive acidic polymer and provide a physical framework for investigating how intracellular polymer mechanics may contribute to auditory mechanotransduction.

## 3 Results

### 3.1 Computational analyses indicate that the PCDH15 CDs are intrinsically disordered

To determine the structural organization of the PCDH15 intracellular region, we first asked whether the three alternatively spliced intracellular isoforms, PCDH15-CD1, PCDH15-CD2 and PCDH15-CD3 (Fig. 1b), are predicted to adopt stable folded conformations. Consensus disorder prediction using Rapid Intrinsic Disorder Analysis Online (RIDAO) indicated that all three CD isoforms are predominantly intrinsically disordered (Fig. 1c)^18^. Although short regions displayed lower disorder scores, none of the isoforms contained a continuous segment predicted to form a stable globular domain.

Sequence composition provided an independent sequence-based indication of disorder. Compared with structured proteins in the Protein Data Bank^19^, all three CDs are depleted in hydrophobic residues and enriched in residues commonly associated with intrinsic disorder, including polar and charged amino acids such as glutamate, serine, threonine and proline (Fig. 1f, Supplementary Fig. 1). Independent AlphaFold3 predictions were consistent with the absence of stable tertiary structure, with low predicted local distance difference (pLDDT) scores throughout most of the cytoplasmic tails (Supplementary Fig. 2)^20^. However, low AlphaFold3 confidence alone cannot distinguish intrinsic disorder from uncertainty in structure prediction^21^.

Although all three isoforms are predicted to be intrinsically disordered, their sequence compositions differ substantially, suggesting that they may exhibit different conformational behaviours. Analysis using the CIDER framework revealed clear differences in the sequence features that are predicted to influence these behaviours (Fig. 1d; Table 1) ^22^. CIDER relates the charge and hydrophobicity of disordered protein sequences to their expected polymer behaviour and maps these sequence properties onto the Das–Pappu phase diagram, which defines regimes associated with different conformational ensembles^22,23^. Three key sequence parameters are the fraction of charged residues (FCR), the net charge per residue (NCPR) and mean hydropathy. FCR describes the overall abundance of charged residues in a sequence, whereas NCPR describes the balance between positive and negative charges. Mean hydropathy describes the average tendency of the amino-acid sequence to favour hydrophobic interactions. Together, these parameters capture important sequence features that influence the conformational behaviour of disordered polymers. PCDH15-CD2 has a substantially higher fraction of charged residues (FCR = 0.39) compared to PCDH15-CD1 (0.19) or PCDH15-CD3 (0.27), placing it within the strong polyampholyte region of the Das–Pappu phase diagram (Fig. 1d; Table 1), whereas PCDH15-CD1 and PCDH15-CD3 occupy the globule/tadpole and Janus boundary regions, respectively^23^. These regimes describe different sequence-encoded tendencies in the conformational behaviour of disordered polymers. The globule/tadpole regime is associated with relatively compact ensembles, whereas the strong polyampholyte regime reflects sequences in which electrostatic interactions are expected to strongly influence chain dimensions. Consistent with this classification, PCDH15-CD2 has a much more negative predicted net charge at neutral pH (−61) than PCDH15-CD1 or PCDH15-CD3 (−1 for each) (Fig. 1e).

**Table 1:** Das–Pappu sequence parameters of the PCDH15 intracellular isoforms. FCR reflects the fraction of charged residues, NCPR the net charge per residue, and mean hydropathy the average hydrophobicity of the sequence. Polymer regimes are assigned according to the Das–Pappu phase diagram.

| Isoform | Length | FCR | NCPR | Mean hydropathy | Predicted polymer regime |
| --- | --- | --- | --- | --- | --- |
| PCDH15-CD1 | 541 | 0.19 | -0.004 | 3.80 | Globule / tadpole |
| PCDH15-CD2 | 381 | 0.39 | -0.16 | 3.26 | Strong polyampholyte |
| PCDH15-CD3 | 280 | 0.27 | -0.004 | 3.71 | Janus boundary |

This unusually high net negative charge arises primarily from the amino acid composition of CD2. PCDH15-CD2 is exceptionally enriched in glutamate residues (24.7%), while simultaneously exhibiting the lowest hydropathy of the three isoforms. In contrast, PCDH15-CD1 is enriched in proline together with serine and threonine residues, and has intermediate charge and hydropathy values (Fig. 1e,f; Supplementary Fig. 1; Table 1).

Together, these analyses suggest that alternative splicing may alter not only the amino acid sequence of the CD, but also its underlying polymer-physical properties. Thus, the three PCDH15 isoforms are predicted to form IDRs with distinct sequence-encoded polymer behaviours rather than simply representing sequence variants of the same polymer. These predictions motivated subsequent biophysical experiments to determine whether the predicted differences in sequence composition translate into distinct conformational properties in solution.

### 3.2 Orthogonal biophysical characterization demonstrates that recombinant CD2 is a predominantly monomeric, expanded IDP

We recombinantly produced and purified two recombinant constructs: (i) CD2, comprising the CD2-specific intracellular region, and (ii) CRCD2, comprising both the common region and the CD2-specific region (Fig. 2a). This allowed us to assess both the intrinsic properties of the CD2-specific region and the contribution of the common region to intracellular protein behaviour. Protein identity was confirmed by mass spectrometry following in-gel digestion, providing independent confirmation that the purified material corresponded to the intended constructs (Supplementary Fig. 3a,b).

**Figure 2.**
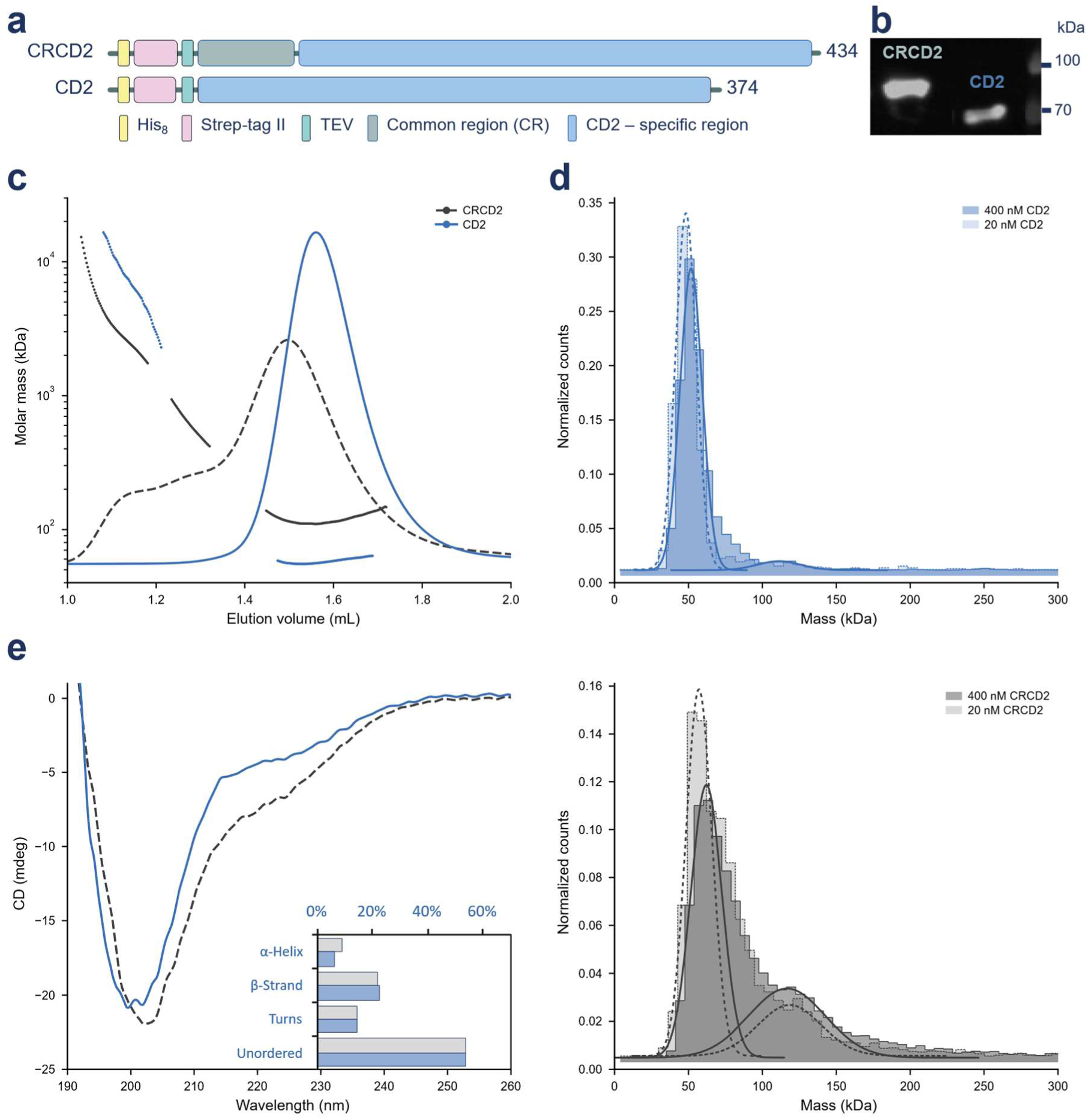
Purification and biophysical characterisation of recombinant PCDH15 intracellular constructs. **(a)** Domain organisation of the recombinant constructs used in this study. CRCD2 contains the conserved common region (CR) fused to the CD2-specific sequence, whereas CD2 contains the CD2-specific intracellular region only. The positions of the His₈ tag, Strep-tag II, TEV cleavage site, CR and CD2-specific region are indicated. **(b)** SDS–PAGE analysis of purified CRCD2 and CD2. Protein identity was independently verified by mass spectrometry following in-gel digestion. **(c)** SEC–MALS profiles of CRCD2 and CD2, showing molar mass as a function of elution volume. **(d)** Mass photometry distributions for CRCD2 and CD2 measured at the indicated concentrations, with fitted molecular-mass populations labelled. **(e)** Far-UV circular dichroism spectra of CRCD2 (grey) and CD2 (blue), with the corresponding estimates of α-helical, β-strand and unordered secondary-structure content shown in the inset.

Across multiple orthogonal techniques, both constructs exhibited solution properties characteristic of IDPs (Fig. 2). SDS–PAGE provided an initial indication of non-globular behaviour, with both constructs displaying anomalous electrophoretic mobility and migrating substantially above their theoretical molecular masses of approximately 42 and 48 kDa, respectively (Fig. 2b). Consistent with their non-globular behaviour, size-exclusion chromatography (SEC) showed markedly earlier elution than expected for globular proteins of similar molecular mass (Supplementary Fig. 3c). Dynamic light scattering further indicated a hydrodynamic radius of approximately 5 nm for CD2, considerably larger than expected for a folded protein of comparable molecular mass (Supplementary Fig. 3d). Together, these measurements indicate that CD2 occupies an unusually large hydrodynamic volume in solution.

Importantly, the enlarged apparent dimensions of CD2 were not attributable to stable oligomerization. Size-exclusion chromatography coupled with multi-angle light scattering (SEC–MALS) measured molecular masses of approximately 49–52 kDa for CD2, close to the theoretical monomeric mass of 42 kDa (Fig. 2c). Mass photometry independently showed that 85–92% of CD2 molecules were monomeric, with only 5–10% detected as dimers (Fig. 2d). Thus, despite its large hydrodynamic dimensions, CD2 behaves predominantly as a monomer in solution, indicating that its early elution from SEC reflects conformational expansion rather than stable oligomerization.

In contrast, CRCD2 showed evidence of substantial self-association, with SEC–MALS detecting a major species of approximately 107 kDa and mass photometry revealing dimer populations of 18–58% depending on the SEC fraction analysed (Fig. 2c,d). This indicates that inclusion of the common region promotes intermolecular association within the intracellular domain.

Far-UV circular dichroism spectroscopy provided an independent assessment of secondary structure. Both constructs exhibited a single minimum at approximately 200 nm, consistent with predominantly disordered conformations lacking substantial α-helical or β-sheet secondary structure (Fig. 2e). Differential scanning fluorimetry did not reveal a defined thermal unfolding transition, further supporting the absence of a stable globular fold (Supplementary Fig. 3e).

Together, these complementary biophysical measurements demonstrate that recombinant CD2 behaves as a predominantly monomeric, expanded disordered protein in solution, in agreement with the sequence-based predictions. In contrast, inclusion of the common region markedly increases intermolecular association, identifying the conserved common region as an important determinant of self-association within the intracellular domain. Having established that the CD2 construct is predominantly monomeric and highly expanded in solution, we next used small-angle X-ray scattering to determine its global dimensions.

### 3.3 SAXS reveals the salt-dependent expansion of the CD2 conformational ensemble

Having established that the recombinant CD2 construct behaves as a predominantly monomeric IDP, we next used small-angle X-ray scattering (SAXS) to determine the properties of its conformational ensemble and test whether its solution behaviour is consistent with the polymer properties predicted from sequence analysis.

SAXS measurements were collected over a six-point concentration series at low (10 mM NaCl) and physiological-like (150 mM NaCl) ionic strength, allowing concentration-dependent effects to be assessed and scattering profiles to be extrapolated to infinite dilution (Fig. 3a,b; Supplementary Fig. 4). At 10 mM NaCl, increasing protein concentration produced systematic low-*q* deviations consistent with repulsive interparticle interactions, whereas concentration-dependent effects were less pronounced at 150 mM NaCl (Supplementary Fig. 4). The two lowest-concentration datasets at 10 mM NaCl showed an additional low-*q* contribution consistent with trace aggregation and were excluded from concentration extrapolation. All six datasets at 150 mM NaCl passed the Guinier quality criteria and were retained for extrapolation. Guinier fits for the retained individual datasets showed appropriate fitting regions without strong systematic deviations in the residuals (Supplementary Fig. 5; Supplementary Table 1).

**Figure 3.**
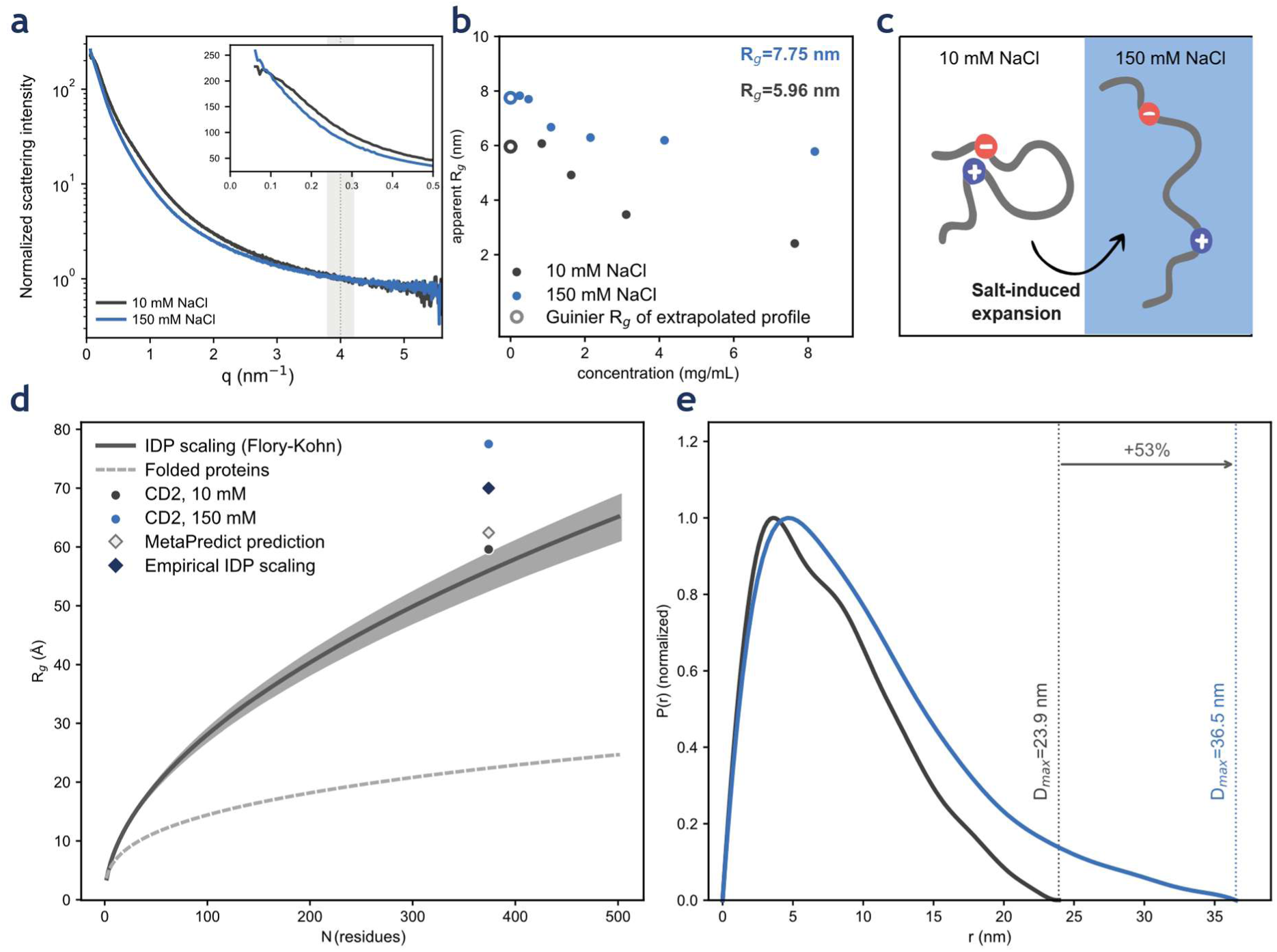
SAXS reveals salt-dependent expansion of the CD2 conformational ensemble. Small-angle X-ray scattering (SAXS) analysis of recombinant CD2 in 10 mM NaCl (dark grey) and 150 mM NaCl (blue). **(a)** Concentration-extrapolated scattering profiles normalized to the forward scattering intensity, I(0), as a function of the scattering vector, q. **(b)** Apparent radius of gyration (Rg) determined by Guinier analysis across the measured protein concentrations. Filled circles show individual measurements; open circles show Guinier Rg values determined from the corresponding concentration-extrapolated scattering profiles. **(c)** Schematic representation of the salt-dependent expansion of the CD2 conformational ensemble. **(d)** Experimental Guinier Rg values from the concentration-extrapolated scattering profiles compared with scaling relationships for folded proteins and intrinsically disordered proteins (IDPs), together with sequence-based MetaPredict and empirical IDP scaling predictions. **(e)** Pair-distance distribution functions, P(r), calculated from the concentration-extrapolated scattering profiles. Vertical dotted lines indicate the maximum particle dimensions (Dmax).

Concentration-extrapolated scattering profiles yielded Guinier radii of gyration (*R*g) of 5.96 nm at 10 mM NaCl and 7.75 nm at 150 mM NaCl. Analysis of the same scattering profiles in real space yielded corresponding pair-distance distribution (*P(r))* derived *R*g values of 6.50 and 8.76 nm and maximum particle dimension (Dmax) values of 23.90 and 36.51 nm, respectively (Supplementary Table 1; Fig. 3a,b,e). Thus, increasing the NaCl concentration increased the Guinier-derived *R*g by approximately 30% and *D*max by approximately 53%.

The measured dimensions are substantially larger than expected for folded/globular proteins of similar mass (Fig. 3d). Sequence-based MetaPredict and empirical IDP scaling predict *R*g values of approximately 6.2 and 5.6 nm, respectively, whereas an independent empirical relationship for IDPs predicts an *R*g of approximately 7.0 nm for CD2 based on its sequence length^24,25^. The experimentally determined *R*g at 10 mM NaCl falls within this range of IDP reference predictions, whereas at 150 mM NaCl it exceeds all three estimates (Fig. 3d). Thus, CD2 is expanded relative to a folded protein under both conditions, with a pronounced additional expansion at higher NaCl concentration.

The increase in dimensions with salt concentration was accompanied by a broad, asymmetric *P(r)* distribution, consistent with an extended conformational ensemble rather than a compact globular structure (Fig. 3e). Dimensionless Kratky analysis likewise supported a disordered, non-compact conformation under both conditions (Supplementary Fig. 5). Increasing salt concentration therefore expanded rather than compacted the CD2 ensemble. This behaviour is consistent with its sequence-predicted polyampholyte character: whereas electrostatic screening generally reduces repulsive interactions in polyelectrolytes, screening of attractive interactions between oppositely charged residues in polyampholytes can favour chain expansion. The observed salt-dependent expansion therefore provides experimental support for the polymer behaviour predicted from the CD2 sequence.

Together, these SAXS measurements demonstrate that CD2 forms an expanded, salt-responsive conformational ensemble whose dimensions increase substantially with NaCl concentration, consistent with its sequence-predicted strong polyampholyte character.

### 3.4 The expanded CD2 intracellular domain contains a conserved signalling hotspot

The absence of stable secondary or tertiary structure does not imply a lack of functional organization. IDPs frequently contain short linear motifs (SLiMs) that mediate interactions with signalling proteins, kinases, phosphatases and cytoskeletal adaptors. Having established that CD2 behaves as an expanded acidic polyampholyte, we next asked whether its sequence also contains conserved regulatory features.

Analysis of the CD2 intracellular domain using the Eukaryotic Linear Motif (ELM) resource identified a dense collection of predicted interaction motifs, several of which are strongly conserved across mammalian orthologues (Fig. 4, Supplementary Fig. 6; Supplementary Table 2). Rather than being distributed uniformly throughout the sequence, many coincide with the glutamate-rich region that contributes to the distinctive sequence composition of CD2.

**Figure 4.**
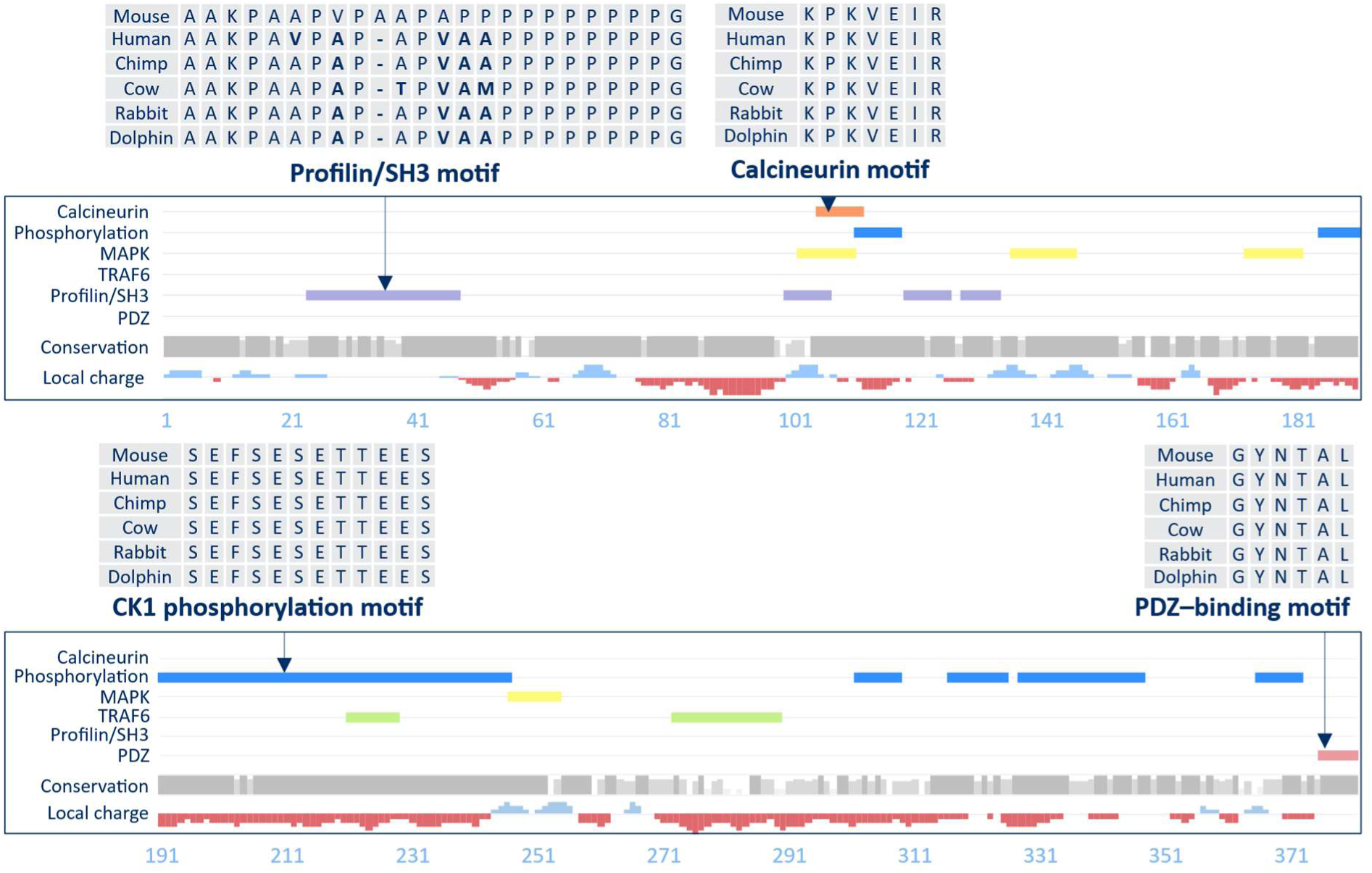
Conserved sequence features and predicted functional motifs within the PCDH15-CD2 intracellular domain. Sequence-based analysis of the PCDH15-CD2 intracellular domain, including the common region, showing predicted short linear motifs, sequence conservation and local charge distribution. Sequence alignments highlight representative conserved features across six mammalian PCDH15-CD2 orthologues, including a proline-rich profilin/SH3-binding region, a calcineurin-binding PxIxIT motif, a representative predicted CK1 phosphorylation motif and the C-terminal class I PDZ-binding motif. Motif tracks show predicted calcineurin, phosphorylation-associated, MAPK, PDZ, profilin/SH3 and TRAF6 sites. Overlapping predictions of the same functional class are merged for visualization. Predicted motifs and their sequence coordinates are listed in Supplementary Table 2.

One of the most prominent features is a PxIxIT motif (KPKVEIR) within the CD2-specific region, matching the canonical docking sequence for the Ca²⁺/calmodulin-dependent phosphatase calcineurin. The motif is invariant across all six mammalian orthologues analysed and, to our knowledge, has not previously been described in PCDH15. Its conservation identifies this sequence as a candidate calcineurin docking site in PCDH15.

The CD2-specific region also contains a pronounced cluster of predicted phosphorylation-associated motifs within the acidic, glutamate-rich region (Fig. 4). Several of these motifs are strongly conserved across mammalian orthologues (Supplementary Table 2). Their overlap with the acidic region raises the possibility that post-translational modification could modulate the conformational properties of CD2.

Additional sequence features include proline-rich Src homology 3 (SH3)/profilin-binding motif predictions, mitogen-activated protein kinase (MAPK) docking motifs, TNF receptor-associated factor 6 (TRAF6)-binding motif predictions and a highly conserved C-terminal class I PDZ-binding motif (Fig. 4). Although these interactions remain to be experimentally validated, the conservation of several prominent motifs identifies them as candidate regulatory features.

Together, these analyses reveal a cluster of candidate regulatory motifs within the expanded CD2 intracellular domain. Their concentration within and around the acidic region identifies a potential signalling hotspot embedded within the sequence features that determine the unusual polymer properties of CD2.

### 3.5 Molecular dynamics simulations reveal transient local interactions within the signalling hotspot

To investigate the local conformational behaviour of the motif-rich region identified by sequence analysis, we performed five independent 1-µs all-atom molecular dynamics (MD) simulations of a 30-residue CD2 peptide spanning an overlapping cluster of predicted SH3-, MAPK-, calcineurin PxIxIT- and CK2-associated motifs (Fig. 5a). The simulated sequence corresponds to residues 93–122 of the CD2 intracellular domain and encompasses the fully conserved PxIxIT motif.

**Figure 5.**
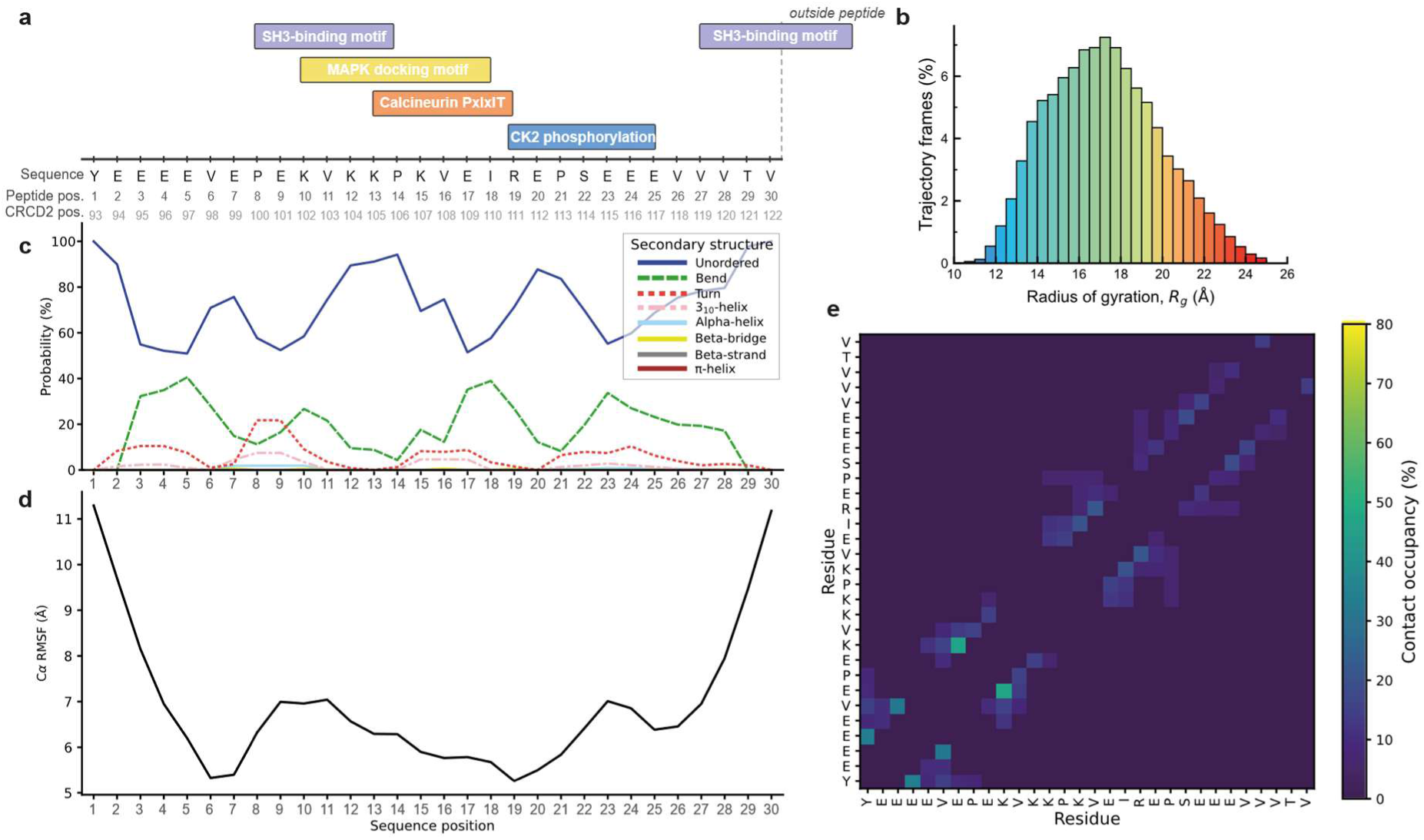
Molecular dynamics reveals a heterogeneous conformational ensemble within a motif-rich region of PCDH15-CD2. **(a)** Predicted regulatory motifs overlapping the 30-residue CD2 peptide used for molecular dynamics simulations (CRCD2 residues 93–122). Motif coordinates are based on Eukaryotic Linear Motif (ELM) predictions; the C-terminal SH3-binding motif extends beyond the simulated sequence. **(b)** Distribution of the radius of gyration (*Rg*) across analysed trajectory frames. **(c)** Per-residue secondary-structure occupancy across the analysed trajectories, showing predominantly unordered conformations with transient local secondary-structure formation. **(d)** Cα root-mean-square fluctuation (RMSF) per residue. **(e)** Residue–residue contact occupancy across the analysed trajectories, illustrating transient local contacts within the peptide. Data represent the concatenation of five independent 1-µs simulations, with the first 300 ns of each trajectory excluded from analysis.

The peptide remained highly dynamic across the simulations, sampling a broad range of conformations and radii of gyration (Fig. 5b), with backbone RMSD trajectories further reflecting substantial conformational variability throughout the simulations (Supplementary Fig. 7a). Secondary-structure analysis of the concatenated trajectories showed predominantly unordered conformations, interspersed with transient bends, turns and short helical segments, with no persistent secondary-structure element (Fig. 5c). Residue-wise fluctuations were non-uniform, with comparatively reduced flexibility across the central motif-overlap region relative to the peptide termini (Fig. 5d).

Despite this conformational heterogeneity, contact analysis of the concatenated trajectories revealed non-uniform transient intramolecular interactions within the peptide (Fig. 5e; Supplementary Fig. 7b). Contacts were predominantly local and distributed across the sequence, including within the region containing the overlapping predicted SH3, MAPK and PxIxIT motifs and the adjacent CK2-associated sequence. These interactions remained transient, with no persistent long-range contact pattern indicative of a stable tertiary structure.

Together, these simulations indicate that the motif-rich CD2 sequence remains conformationally dynamic while supporting transient local interactions within the overlapping regulatory region. Thus, rather than forming a stable structural element, this conserved sequence retains substantial conformational heterogeneity while exhibiting local conformational organization.

## 4 Discussion

The present study identifies the cytoplasmic domain (CD) of the hearing-essential PCDH15-CD2 isoform as a highly charged, environmentally responsive IDR with unusual polymer properties. Sequence analysis, orthogonal biophysical characterisation and SAXS show that CD2 lacks stable tertiary structure, adopts an expanded conformational ensemble and undergoes substantial salt-dependent changes in its dimensions. In parallel, sequence analysis revealed a conserved motif-rich region containing candidate regulatory sites, including an invariant calcineurin-binding PxIxIT motif. Together, these findings suggest that the PCDH15 CD should not be considered solely as a passive molecular tether, but as a dynamic polymer whose physical and regulatory properties may contribute to the organisation and function of the mechanotransduction complex.

Current models of auditory mechanotransduction have largely focused on force transmission through the extracellular tip link and the membrane-associated MET machinery^1,5^. The extracellular cadherin repeats of PCDH15 and CDH23 form the mechanically loaded tip-link filament^2,9^, while TMC1, TMIE, CIB2 and associated proteins form part of the membrane-associated MET complex at its lower insertion site^3,5–8^. By comparison, the potential mechanical contribution of the large PCDH15 CD remains poorly understood. Our findings provide a physical basis for reconsidering this region. Importantly, they do not establish CD2 as the intracellular elastic element or gating spring, nor do they demonstrate force transmission by CD2 in hair cells^26^. Rather, they identify sequence-encoded polymer properties that could influence how force is transmitted or regulated on the intracellular side of PCDH15.

The pronounced salt sensitivity of CD2 provides a potential mechanism through which its intracellular environment could influence its conformational ensemble. Increasing NaCl from 10 to 150 mM substantially expanded the isolated CD2 chain, consistent with polyampholyte behaviour in which electrostatic screening weakens attractive interactions between oppositely charged residues^23,27–29^. The consequences may be different, but complementary, when these properties are considered in the native architecture of PCDH15. PCDH15 forms a cis-dimer, thereby positioning the membrane-proximal origins of its two intracellular tails in close proximity^10,30–32^. Because the CD2 tails carry a strong net negative charge, this geometry creates the possibility of electrostatic interactions not present in the isolated-chain SAXS measurements. In this paired geometry, increased electrostatic screening could affect intra- and interchain interactions differently. Within each CD2 chain, screening could weaken attractive interactions and favour the expansion observed experimentally. At the same time, screening could reduce repulsion between the two negatively charged tails, allowing their conformational ensembles to overlap more extensively. Changes in this balance between intrachain and interchain electrostatic interactions could therefore reorganize the paired CD2 ensemble and, if mechanically constrained, alter its extension or mechanical state.

The common region may provide an additional level of organization to such a paired system. Whereas the isolated CD2-specific construct remained predominantly monomeric, inclusion of the common region substantially increased self-association. Although the molecular basis of this association remains unknown, it raises the possibility that the common region constrains the relative organization of the two intracellular domains close to their membrane-proximal origin. Together with interactions of the PCDH15 intracellular region with other components of the hair-cell machinery, including TMC proteins^33^, this self-association could provide constraints within which the electrostatic properties of the CD2 tails become mechanically relevant. The paired-polymer behaviour proposed here nevertheless remains hypothetical, as the present SAXS measurements describe isolated, predominantly monomeric CD2 in solution rather than membrane-anchored pairs under mechanical load.

The regulatory architecture embedded within CD2 provides a second potential means of tuning this ensemble. The CD2-specific region contains an overlapping cluster of predicted SH3-, MAPK-, calcineurin PxIxIT- and phosphorylation-associated motifs, including an invariant PxIxIT sequence adjacent to a conserved phosphorylation-rich acidic region containing predicted CK2 recognition sites. MD simulations of a 30-residue peptide spanning this motif-rich region showed that the sequence remained conformationally heterogeneous while forming transient local contacts within the overlapping motif cluster. These simulations therefore suggest that local conformational organisation can emerge without formation of a stable fold, consistent with the broader behaviour of IDRs in which transient local structure and intramolecular contacts can influence molecular recognition^34^. The extensive overlap between predicted motifs is particularly intriguing because recognition of one site could potentially alter the accessibility or local context of neighbouring motifs, enabling context-dependent interactions within the disordered tail^35,36^. The simulations were restricted to a short isolated peptide and do not establish any of the predicted protein interactions, but provide a structural basis for investigating how local sequence dynamics and motif accessibility may be coupled within CD2.

The invariant PxIxIT motif is particularly notable in this context because PxIxIT sequences mediate docking to the Ca²⁺/calmodulin-dependent phosphatase calcineurin^37^. Its proximity to a phosphorylation-rich acidic region places a candidate Ca²⁺-dependent phosphatase docking site alongside a potential phosphoregulatory region. Phosphorylation can alter the charge patterning and intramolecular interactions of IDRs, thereby reshaping their local and global conformational ensembles^38,39^. Whether calcineurin binds PCDH15, whether the predicted Ser/Thr sites are phosphorylated in hair cells, and how their phosphorylation state affects CD2 remain unknown. Nevertheless, the juxtaposition of a conserved calcineurin-docking motif, phosphorylation-associated sequences and a dynamic overlapping motif cluster provides a testable connection between biochemical signalling and the unusual polymer properties of CD2.

Together, these observations suggest a two-timescale mechanochemical model in which the paired CD2 tails respond to rapid changes in electrostatic conditions and slower biochemical regulation (Fig. 6). At rest, the two highly charged tails are proposed to form a dynamic ensemble between their membrane attachment sites and an intracellular anchoring network. On a rapid timescale, tip-link tension mechanically loads the paired CD2 tails, with electrostatic repulsion between the negatively charged chains opposing their deformation. MET-channel opening then produces an inward current carried predominantly by K⁺, with a smaller Ca²⁺ component^40^. We hypothesize that the local ionic flux associated with channel opening could transiently alter electrostatic screening around the CD2 tails. Within individual chains, increased screening could weaken attractive interactions between oppositely charged residues, favouring the expansion observed experimentally at higher ionic strength. Screening could also reduce repulsive interactions between the predominantly negatively charged tails, permitting greater overlap of their conformational ensembles. In the constrained geometry of a membrane- and intracellularly anchored pair, this redistribution could rapidly alter the extension and tension of the ensemble without requiring a defined structural transition or enzymatic modification. The SAXS measurements establish salt responsiveness of isolated CD2 but do not demonstrate that MET-channel opening produces a sufficient local ionic perturbation, or that the resulting conformational changes alter force transmission in vivo.

**Figure 6.**
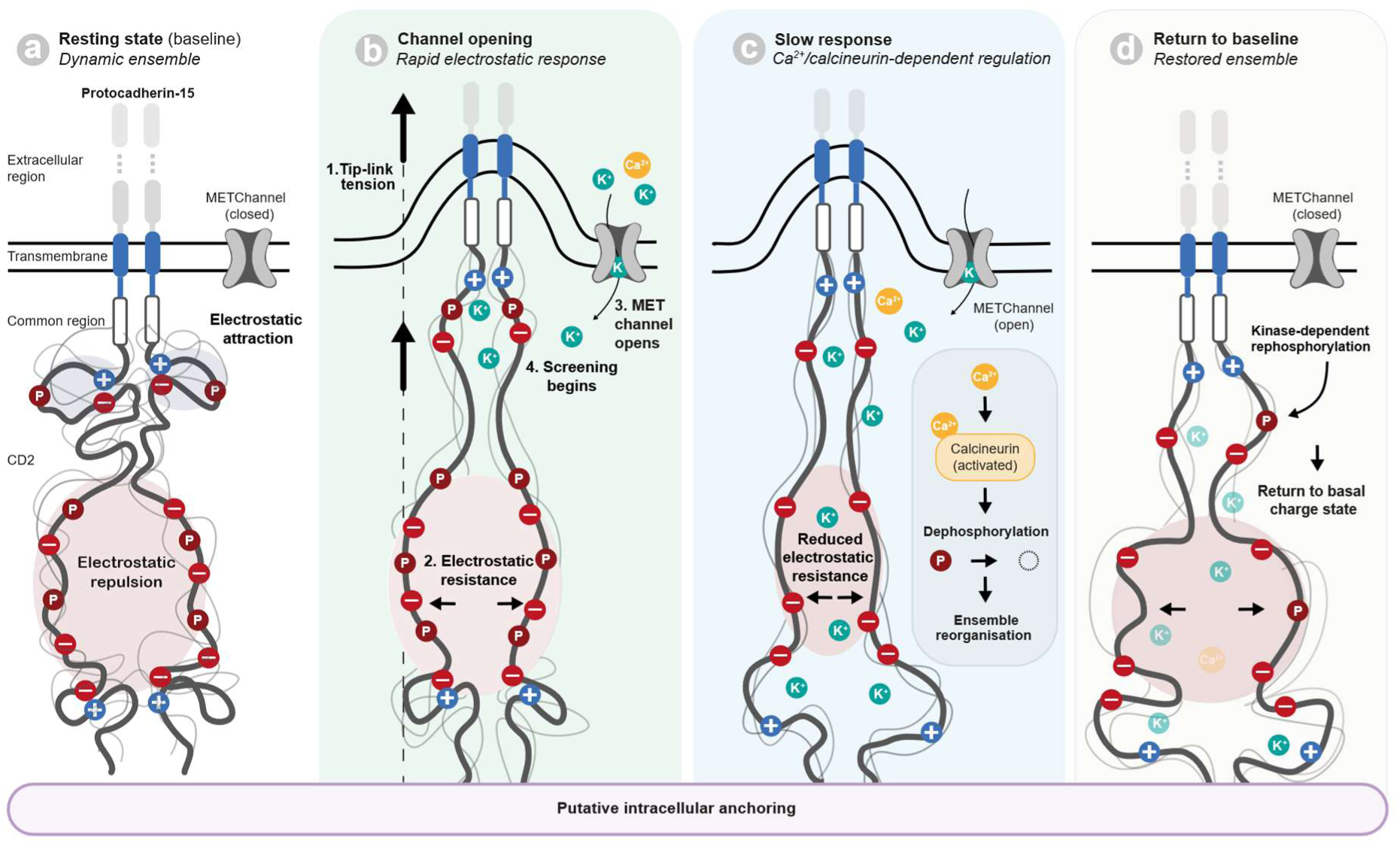
Proposed two-timescale mechanochemical model for regulation of the PCDH15-CD2 cytoplasmic ensemble. Schematic illustrating a hypothetical model in which the highly charged, intrinsically disordered PCDH15-CD2 tails form a dynamically regulated paired ensemble. **(a)** At rest, the paired tails adopt a dynamic ensemble shaped by attractive and repulsive intra- and interchain electrostatic interactions. **(b)** Tip-link tension loads the paired CD2 tails, with electrostatic repulsion between the negatively charged tails opposing their deformation. MET-channel opening then initiates ion influx, which could transiently alter electrostatic screening around the CD2 tails. **(c)** Electrostatic screening could reduce interchain repulsion while altering intrachain interactions, reorganizing the paired ensemble and reducing its electrostatic resistance. In parallel, Ca²⁺ entry could activate calcineurin through Ca²⁺/calmodulin-dependent signalling and facilitate its recruitment through the conserved PxIxIT motif. Subsequent dephosphorylation of the neighbouring phosphoregulatory region could reduce negative charge and further modify the electrostatic, conformational and mechanical state of the tails. **(d)** Kinase-dependent rephosphorylation could restore the basal charge distribution and favour return towards the resting ensemble.

A slower regulatory process could operate through Ca²⁺-dependent signalling. During sustained MET-channel activity, Ca²⁺ entry through the MET channel could activate calcineurin through Ca²⁺/calmodulin-dependent signalling and facilitate its recruitment to CD2 through the conserved PxIxIT motif^40,41^. We hypothesize that calcineurin recruitment could promote dephosphorylation within the neighbouring phosphoregulatory region, thereby reducing negative charge within this region and altering the electrostatic interaction landscape of the paired tails, which could further modify their conformational and mechanical state. The local conformational organisation observed in the MD simulations further suggests that changes in charge or binding within this region could influence the context and accessibility of overlapping motifs. Subsequent kinase-dependent rephosphorylation could provide a route towards restoration of the basal ensemble. In this framework, rapid electrostatic screening and slower post-translational regulation act on the same disordered polymer over distinct timescales, providing a potential mechanism for coupling immediate electrostatic responses to longer-term regulation of the intracellular ensemble.

This model is intended to complement rather than replace existing mechanisms of mechanotransduction and adaptation. Hair-cell adaptation encompasses processes operating across different timescales, whose molecular basis and Ca²⁺ dependence remain debated^5,42^. The proposed mechanism therefore does not assign electrostatic or phosphorylation-dependent regulation of CD2 to any single established component of adaptation. Instead, it introduces the PCDH15 intracellular ensemble as an additional physical layer through which changes in mechanical state could arise from redistribution of a disordered conformational ensemble rather than from a defined folding transition. We propose that such regulation could contribute to mechanotransduction on different timescales without equating the two proposed processes directly with established fast and slow adaptation mechanisms.

Several aspects of this framework remain to be established. Our biophysical measurements were performed on isolated recombinant intracellular constructs in solution and therefore do not reproduce the paired geometry, geometric constraints, molecular crowding, membrane attachment or binding network encountered by CD2 in stereocilia. The MD simulations examine only a 30-residue segment and cannot establish the behaviour of the full CD2 ensemble. Moreover, neither calcineurin binding nor phosphorylation of the predicted sites has been demonstrated, and we have not directly measured CD2 elasticity or force transmission. Key tests of the model will therefore include determining whether MET-channel activity measurably alters the local electrostatic environment of CD2, identifying its interaction partners and phosphorylation state in hair cells, measuring how phosphorylation affects its conformational and mechanical properties, and examining paired CD2 tails in membrane-associated systems under mechanical load. Perturbation of the PxIxIT motif and phosphoregulatory region in hair cells would provide a direct means of testing whether the proposed regulatory architecture contributes to mechanotransduction or adaptation.

More broadly, these findings suggest that the functional information encoded by the CD2 sequence may reside not in a stable three-dimensional structure, but in the properties of its conformational ensemble. Alternative splicing of PCDH15 generates intracellular isoforms with distinct sequence composition and predicted polymer properties while retaining the same extracellular and transmembrane architecture^11^. In CD2, charge patterning, conformational dynamics and embedded regulatory motifs coexist within the same disordered sequence. These features suggest that CD2 function may emerge from a dynamic ensemble whose physical properties and molecular interactions can be modulated by its environment and regulatory state. Whether this state contributes directly to auditory force transmission remains to be established, but our findings provide a framework for incorporating the intracellular PCDH15 ensemble into models of auditory mechanotransduction and for testing how polymer physics, biochemical regulation and mechanical force intersect within the tip-link complex.

## 5 Materials and methods

### 5.1 Constructs

Two recombinant mouse PCDH15 intracellular constructs were generated: CD2 (Q99PJ1-10; residues 1470–1790; 321 amino acids; theoretical molecular weight 36.5 kDa; theoretical pI 4.19) and CRCD2 (Q99PJ1-10; residues 1410–1790; 381 amino acids; theoretical molecular weight 42.8 kDa; theoretical pI 4.30). Coding sequences were synthesized (GenScript) and cloned into pET-21b(+) as N-terminal fusions to an octahistidine–double Strep-II affinity tag separated from the construct by a TEV protease cleavage site. All biophysical experiments were performed using the tagged proteins. TEV cleavage was not performed because removal of the affinity tag was not required for the experiments described here and reduced the recovery of the highly acidic constructs. Sequence-based analyses were performed on the untagged mature intracellular sequences.

### 5.2 Protein expression and purification

Recombinant constructs were expressed in *Escherichia coli* BL21(DE3). Cells were grown in 2×TY medium at 37 °C to an OD₆₀₀ of approximately 0.6-0.8, induced with 0.05 mM IPTG, and incubated overnight at 18 °C before harvesting by centrifugation.

Cell pellets were resuspended in lysis buffer (20 mM Tris-HCl, 500 mM NaCl, pH 8.0) supplemented with EDTA-free cOmplete protease inhibitor (Roche) and lysed by sonication. Lysates were clarified by centrifugation followed by filtration through a 0.45 μm membrane.

Soluble protein was purified by immobilized metal affinity chromatography using a HisTrap column (Cytiva). After washing with equilibration buffer (20 mM Tris-HCl, 500 mM NaCl, 20 mM imidazole, pH 8.0), bound protein was eluted with a linear gradient of 20–500 mM imidazole. Peak fractions were pooled, concentrated, and centrifuged immediately before SEC to remove insoluble material. Final purification was performed on a HiLoad 16/600 Superdex 200 pg column (Cytiva) equilibrated in 20 mM Tris-HCl, 150 mM NaCl, pH 8.0.

Purified proteins were flash-frozen in liquid nitrogen and stored at −80 °C. Protein concentrations were determined spectrophotometrically at 280 nm using theoretical extinction coefficients calculated from the tagged protein sequences.

### 5.3 SDS–PAGE analysis

Protein purity was assessed by SDS–PAGE under reducing conditions using 7.5% polyacrylamide gels. Proteins were visualized by Coomassie Brilliant Blue staining. A PageRuler Prestained Protein Ladder (Thermo Scientific) was used as the molecular weight standard.

### 5.4 Gel LC/MS

Protein identity was confirmed by liquid chromatography–tandem mass spectrometry (LC–MS/MS). Protein bands were excised from Coomassie-stained SDS–PAGE gels and subjected to in-gel tryptic digestion. Peptides were analysed at the Centre for Proteomics (University of Antwerp) on an Evosep One system (Whisper Zoom 20 SPD) with an Aurora Elite CSI column (15 cm × 75 µm) coupled to a timsTOF SCP mass spectrometer (Bruker Daltonics), operated in data-dependent acquisition mode using parallel accumulation–serial fragmentation (DDA-PASEF). Precursor ions were scanned over an m/z range of 100–1700 with an ion mobility range of 0.80–1.66 V·s·cm⁻² (1/K₀); each PASEF cycle comprised one MS1 scan followed by 10 MS2 ramps (cycle time 1.9 s). Precursors with charge states 1–5 and intensities above 500 were selected for fragmentation, with collision energy ramped from 20 to 59 eV, and active exclusion was enabled with a release time of 0.4 min.

Raw data were processed using FragPipe (v23.1) and searched with MSFragger (v4.3) against the *Mus musculus* database (UP000000589). Trypsin/P was specified as the proteolytic enzyme, allowing up to two missed cleavages. Carbamidomethylation of cysteine was set as a fixed modification, and methionine oxidation and protein N-terminal acetylation as variable modifications. Precursor and fragment mass tolerances were set to ±20 ppm. Peptide-spectrum matches were rescored using Percolator with MSBooster-derived predictions, and false discovery rates were controlled at 1% at the peptide-spectrum match, peptide and protein levels using a target–decoy strategy.

### 5.5 Nano-differential scanning fluorimetry (nanoDSF)

Thermal unfolding profiles of CD2 and CRCD2 were measured using a Prometheus NT.48 nanoDSF system (NanoTemper Technologies GmbH). Protein samples (10 µL; 0.1–0.5 mg mL⁻¹) in 20 mM Tris-HCl, 150 mM NaCl, pH 8.0 were loaded into standard-grade capillaries (NanoTemper Technologies GmbH). Intrinsic protein fluorescence was monitored continuously at 330 and 350 nm while the temperature was increased from 20 to 95 °C at a rate of 1.0 °C min⁻¹ using 40% excitation power. The fluorescence intensity ratio (F350/F330) was plotted as a function of temperature using data exported from PR.ThermControl software (NanoTemper Technologies GmbH).

### 5.6 Circular dichroism (CD) spectroscopy

Far-UV circular dichroism (CD) spectra were recorded at 20 °C using a J-815 spectropolarimeter (Jasco). Protein samples were diluted to 0.5 mg mL⁻¹ in 20 mM Tris-HCl, 150 mM NaCl, pH 8.0. Spectra were collected from 260 to 190 nm at a scan rate of 50 nm min⁻¹ using a bandwidth of 1 nm, a digital integration time of 1 s, and 10 accumulations. A buffer spectrum recorded under identical conditions was subtracted from each protein spectrum.

Secondary-structure content was estimated using the CDSSTR, CONTIN-LL, and SELCON3 algorithms implemented in the DichroWeb server using reference set 7^43–47^. The reported secondary-structure fractions represent the average of the three analyses. A theoretical CD spectrum was calculated from the AlphaFold structural model using PDBMD2CD^48^.

### 5.7 Dynamic Light Scattering

Dynamic light scattering measurements were performed at 20 °C using a Zetasizer mV instrument (Malvern Panalytical). Protein samples (0.1–0.5 mg mL⁻¹) were measured in 20 mM Tris-HCl, 150 mM NaCl, pH 8.0. Two consecutive measurements were collected for each sample using the instrument’s automatic acquisition settings. Data were analysed using the protein analysis model implemented in the Zetasizer software (Malvern Panalytical) to determine the hydrodynamic radius and sample polydispersity.

### 5.8 Mass Photometry

Mass photometry measurements were performed at room temperature using a TwoMP mass photometer (Refeyn Ltd). Protein samples were diluted to 400 nM in 20 mM Tris-HCl, 150 mM NaCl, pH 8.0 and immediately before measurement were further diluted to a final concentration of 20 nM in the same buffer. Measurements were performed on cleaned glass coverslips according to the manufacturer’s recommendations. Videos of 60 s were acquired using AcquireMP (version 2024 R1; Refeyn Ltd) and analysed using DiscoverMP (version 2024 R1; Refeyn Ltd). Contrast-to-mass calibration was performed using bovine serum albumin (BSA) and immunoglobulin G (IgG) standards.

### 5.9 Size-exclusion chromatography coupled with multi-angle light scattering (SEC–MALS)

Protein samples (50 µL, 0.5–1.0 mg mL⁻¹) were injected onto a Superdex 200 Increase 5/150 GL column (Cytiva) equilibrated in 20 mM Tris-HCl, 150 mM NaCl, pH 8.0 using an Agilent 1260 Infinity II HPLC system (Agilent Technologies). Chromatography was performed at room temperature with a flow rate of 0.3 mL min⁻¹. The column was coupled in-line to a MiniDAWN multi-angle light scattering detector and an Optilab differential refractive index detector (Wyatt Technology). Data were analysed using ASTRA version 8.2.2 (Wyatt Technology) using a protein refractive index increment (*dn/dc*) of 0.185 mL g⁻¹.

### 5.10 Small-angle X-Ray scattering (SAXS)

Small-angle X-ray scattering (SAXS) measurements were performed at beamline BM29 of the European Synchrotron Radiation Facility (ESRF, Grenoble, France). Recombinant CD2 was analysed in batch mode over concentration series spanning 0.20–7.64 mg mL⁻¹ in low-salt buffer (20 mM Tris-HCl, 10 mM NaCl, pH 8.0) and 0.25–8.18 mg mL⁻¹ in physiological-like ionic strength buffer (20 mM Tris-HCl, 150 mM NaCl, pH 8.0). Data were collected at 12.5 keV (λ = 0.992 Å) at 20 °C over a q-range of 0.00776–0.495 Å⁻¹. Ten frames with an exposure time of 1 s per frame were collected for each sample. Unless otherwise stated, SAXS data processing and analysis were performed using ATSAS version 3.2.1.

Individual exposure frames were inspected for radiation damage by comparing successive scattering curves prior to averaging. Frames exhibiting radiation-induced changes in the low-*q* region were discarded. Following buffer subtraction, the quality of background subtraction was verified by confirming a flat, symmetric distribution of residual noise around zero at high *q*.

Radius of gyration (*Rg*) values were determined by Guinier analysis using PRIMUS (ATSAS v3.2.1). Because CD2 is an expanded intrinsically disordered protein rather than a compact globular particle, Guinier fitting employed more stringent criteria than the conventional *qRg* < 1.3 limit typically used for globular particles. Acceptable fits were required to satisfy (*i*) *q*maxRg ≲ 1.0–1.3, (*ii*) randomly distributed residuals without systematic deviation, and (*iii*) stability of the calculated *Rg* following small adjustments to the fitting window. The selected fitting ranges were independently verified using the automated fitting routine AUTORG, which yielded essentially identical *Rg* values.

For the two lowest-concentration datasets collected under low-salt conditions (0.20 and 0.44 mg mL⁻¹), inclusion of the lowest-*q* data produced a systematic increase in apparent *Rg*, suggesting a concentration-independent trace aggregate population. These two datasets were therefore excluded from the concentration extrapolation used for structural analysis, leaving four concentrations (0.84–7.64 mg mL⁻¹) for the low-salt condition. All six concentrations were retained for the physiological-salt dataset.

For the remaining low-salt measurements, increasing protein concentration produced systematic low-*q* deviations characteristic of repulsive interparticle interactions. To minimise these effects, fitting windows were selected to reduce the contribution of the affected low-*q* region while retaining at least six data points for stable linear regression.

To account for residual concentration-dependent effects, *R*g, *I*(0)/c, and the scattering intensity at each *q* value were extrapolated to infinite dilution by linear regression as a function of protein concentration using the extrapolation routines implemented in ATSAS. The concentration dependence of *I*(0)/c was used as a qualitative indicator of intermolecular interactions.

Pair-distance distribution functions (*P(r)*) and maximum particle dimensions (*D*max) were calculated in GNOM from the concentration-extrapolated scattering curves. *D*max was optimized iteratively to obtain smooth *P(r)* distributions that approached zero at the maximum intramolecular distance. *R*g values obtained from the *P(r)* distributions were compared with Guinier-derived values as an internal consistency check.

Dimensionless Kratky plots were calculated from the concentration-extrapolated scattering curves to assess the global conformational state of CD2. Statistical differences between scattering profiles were evaluated using the CorMap test as implemented in ATSAS.

### 5.11 Molecular dynamics simulations

All-atom MD simulations were performed using GROMACS 2023 to investigate the CD2 calcineurin PxIxIT motif-containing peptide in aqueous solution containing 10 mM NaCl. The peptide was described using the AMBER99SB-ILDN force field, while water molecules were represented using the TIP4P-D model. The initial structure was generated as an extended linear chain using Avogadro version 1.2.0. Standard protonation states at neutral pH and charged N-and C-termini were used, giving the peptide a net charge of -6e. The peptide was placed in a dodecahedral simulation box with periodic boundary conditions, maintaining a minimum distance of 1.0 nm between the peptide and the box boundaries. The system was solvated, and water molecules were replaced with Na⁺ and Cl⁻ ions to obtain a NaCl concentration of 10 mM while neutralizing the total system charge. The equations of motion were integrated using the leapfrog algorithm with a time step of 2 fs. Short-range nonbonded interactions were treated using the Verlet cutoff scheme with a cutoff of 1.2 nm. Long-range electrostatic interactions were calculated using the particle mesh Ewald method with cubic interpolation and a Fourier grid spacing of 0.16 nm. Long-range dispersion corrections were applied to both the energy and pressure. All bonds involving hydrogen atoms were constrained using the LINCS algorithm. The temperature was maintained at 298 K using the Nosé–Hoover thermostat with a coupling time constant of 1.0 ps, with the peptide coupled separately from the remaining system. Isotropic pressure coupling was applied using the Parrinello–Rahman barostat with a reference pressure of 1 bar, a coupling time constant of 5.0 ps, and a compressibility of 4.5 × 10⁻⁵ bar⁻¹. Energy minimization was performed using the steepest-descent algorithm. This was followed by three equilibration stages: 0.5 ns in the NVT ensemble, 0.5 ns in the NPT ensemble, and a further 1.0 ns in the NPT ensemble. Position restraints were applied to the peptide during all equilibration stages. Five independent production simulations of 1 μs each were subsequently performed, giving a total simulation time of 5 μs.

### 5.12 Trajectory analysis

The first 300 ns of each independent simulation replicate were excluded from the analysis to minimize the influence of the initial relaxation and conformational adaptation of the extended starting peptide. The remaining 700 ns from each of the five independent simulations were concatenated using the GROMACS gmx trjcat tool, resulting in a combined analysed trajectory of 3.5 μs containing 350,000 frames.

The conformational dimensions of the peptide were characterized by calculating the radius of gyration (Rg) using the GROMACS gmx gyrate tool. The trajectory frames were subsequently classified according to their Rg values using an in-house Python script. The script generated frame-index files compatible with GROMACS, which were used together with gmx trjconv to extract separate trajectories corresponding to different Rg-based conformational groups. These subsets were subsequently used for further structural analyses.

The root-mean-square deviation (RMSD) was calculated using the GROMACS gmx rms tool. The distribution of RMSD values sampled throughout the combined trajectory was visualized using kernel density estimation (KDE) implemented with the kdeplot function of the Seaborn Python library, providing a continuous representation of the probability density of the sampled RMSD values.

Residue-specific conformational flexibility was quantified by calculating the root-mean-square fluctuation (RMSF) of the Cα atom of each residue over the analysed trajectory. Prior to calculation of the RMSF, overall translational and rotational motions of the peptide were removed by structural alignment of the trajectory. The resulting Cα RMSF profile was used to identify comparatively flexible and conformationally restricted regions along the peptide sequence.

Secondary-structure formation was analysed on a residue-by-residue and frame-by-frame basis using the DSSP secondary-structure classification. The secondary-structure states were retained using the full DSSP classification, corresponding to unordered, bend, turn, 3₁₀-helix, α-helix, β-bridge, β-strand, and π-helix. For each residue and each secondary-structure class, the occupancy was calculated as the percentage of analysed frames in which that residue was assigned to the corresponding secondary-structure state. The resulting occupancies therefore represent the fraction of the simulated ensemble in which each residue adopted a particular secondary-structure state.

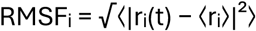

where rᵢ(t) represents the position of the Cα atom of residue i at time t and ⟨rᵢ⟩ is its time-averaged position.

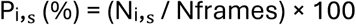

where Nᵢ,ₛ is the number of analysed frames in which residue i was assigned to secondary-structure state s and Nframes is the total number of analysed trajectory frames.

Intramolecular residue–residue contacts were analysed throughout the combined trajectory to characterize transient interactions within the peptide. Two residues were considered to form a contact when at least one pair of atoms belonging to the two residues was separated by a distance of no more than 8 Å. Contacts between residues close in primary sequence were excluded by omitting residue pairs with a sequence separation of |i − j| < 3, thereby reducing contributions from trivial local interactions associated with covalent connectivity and immediate sequence neighbours.

For each residue pair i,j, contact occupancy was calculated as:

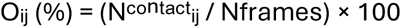

where Nᶜᵒⁿᵗᵃᶜᵗᵢⱼ is the number of analysed frames in which residues i and j fulfilled the contact criterion. The resulting residue–residue contact-occupancy matrix was used to identify recurrent and transient intramolecular interactions sampled by the peptide ensemble. Contact occupancies were represented as percentages in the two-dimensional residue contact map.

Interresidue distances, backbone dihedral-angle distributions, and hydrogen-bond interactions were analysed using the GROMACS gmx pairdist, gmx rama, and gmx hbond tools, respectively. Ramachandran distributions were represented as two-dimensional histograms of the backbone φ and ψ dihedral angles, allowing the populations of different regions of conformational space to be quantified.

Finally, structural clustering was performed using the GROMOS clustering algorithm implemented in the GROMACS gmx cluster tool. An RMSD cutoff of 0.9 nm was used to group structurally similar conformations, thereby enabling identification of representative conformational states sampled over the combined trajectory.

### 5.13 Bioinformatic analysis

#### 5.13.1 Disorder prediction

Sequence-based analyses were performed using the mouse PCDH15-CD1 (UniProt Q99PJ1-1), PCDH15-CD2 (Q99PJ1-10) and PCDH15-CD3 (Q99PJ1-18) isoforms. Intrinsic disorder was predicted for the intracellular regions of all three PCDH15 isoforms using the RIDAO consensus meta-predictor, which integrates predictions from PONDR VLXT, PONDR VL3, PONDR VSL2, PONDR FIT, and IUPred (short and long)^18^. Unless otherwise indicated, disorder analyses refer to the untagged mature intracellular sequences. Predicted radii of gyration were obtained using the MetaPredict web server on tagged CD2^49^.

#### 5.13.2 Sequence-composition analysis

Sequence-composition descriptors including the fraction of charged residues (FCR), net charge per residue (NCPR), mean hydropathy, charge segregation (κ), Ω, and Das–Pappu phase behaviour classification were calculated using CIDER^22^. Amino acid composition, molecular weight, theoretical isoelectric point, extinction coefficient, and related physicochemical properties were calculated using ProtParam. To contextualize amino acid composition within known protein classes, per-residue frequencies of CD1–CD3 were compared against the reference composition of structured proteins in the PDB Select 25 dataset, as reported by Composition Profiler^19^.

#### 5.13.3 AlphaFold predictions

Amino acid sequences of the CD1–CD3 cytoplasmic tails were submitted as single protein chains to the AlphaFold Server (accessed 27 November 2024), which implements AlphaFold3. For each sequence, five models were generated under default server settings (single random seed, 10 recycles), and the top-ranked model was used for analysis. Model confidence was assessed using the per-residue predicted local distance difference test, with pLDDT < 50 taken as indicative of low model confidence. AlphaFold3 additionally reports a per-target fraction of disordered residues, which was 0.99, 1.00 and 1.00 for CD1, CD2 and CD3, respectively.

#### 5.13.4 Short linear motifs

The intracellular region of each isoform was scanned against the ELM regular-expression library^50^, restricted to cytoplasmic-compartment ELM classes. Candidate motifs were considered conserved when the complete motif sequence was present in all six mammalian orthologues analysed.

#### 5.13.5 Cross-species alignment

Intracellular CD2 sequences from *Mus musculus* (UniProt Q99PJ1-10), *Homo sapiens* (Q96QU1-4), *Pan troglodytes* (A0A2I3T0A9), *Bos taurus* (G3MYD4), *Oryctolagus cuniculus* (G1TZM4) and *Tursiops truncatus* (A0A2U4BFX7) were aligned using MAFFT v7.511 with the L-INS-i algorithm.

## 6 Data availability

The SAXS data generated in this study have been deposited in the Small Angle Scattering Biological Data Bank (SASBDB) under accession codes SASD2B8 (10 mM NaCl) and SASD2C8 (150 mM NaCl). The molecular dynamics simulation data generated in this study are available through Zenodo at <u>10.5281/zenodo.22745618</u>.

## 7 Code availability

The custom Python code used for analysis of the molecular dynamics simulations is available through Zenodo at 10.5281/zenodo.22745618.

## 9 Author Contributions

J.V.d.V., Y.S., M.S., P.V.W., and V.V.R. contributed to study conceptualisation. J.V.d.V. designed and performed the sequence-based computational analyses and biophysical experiments and analysed the data. K.L. contributed to the biophysical experiments. M.S. supervised the development of the SAXS studies and contributed resources and methodological expertise. N.M. performed SAXS data collection and contributed to SAXS data analysis. N.M. developed and performed the molecular dynamics simulations, with supervision and resources provided by M.S.; N.M. and M.S. analysed and interpreted the simulation data. R.B. performed the LC–MS experiments and generated the proteomic data that motivated this study. J.V.d.V., M.S., Y.S., P.V.W., V.V.R., and C.P. contributed to interpretation of the results. J.V.d.V. drafted the manuscript. R.B., K.L., N.M., M.S., Y.S., P.V.W., V.V.R., and C.P. provided critical comments and revised the manuscript. M.S., V.V.R., and C.P. acquired funding. All authors approved the final manuscript.

## 10 Competing interests

The authors declare no competing interests.

## 11 Funding

This work was financially supported by the Research Foundation – Flanders (FWO; Project G052622N and Senior Clinical Investigator Grant 18E2524N to V.V.R.). Additional support was provided through an FWO Travel Grant (K243925N) and an EMBL Christian Boulin Fellowship. This research was funded by a grant from Fondation pour l’Audition (FPA-IDA05 to C.P). This work also benefited from a French government grant managed by the Agence Nationale de la Recherche under the France 2030 program, reference ANR-23-IAHU-0003.

## 12 Acknowledgments

We thank the EMBL Heidelberg Protein Expression and Purification Core Facility, in particular Kim Remans, for access to the facility. We thank the Division of Computational Chemistry for facilitating and hosting part of this work at Lund University.

We acknowledge the simulation resources provided by the National Academic Infrastructure for Supercomputing in Sweden (NAISS), which is partially funded by the Swedish Research Council through grant agreement no. 2022-06725 and resources provided by the Swedish National Infrastructure for Computing, SNIC, at the Center for Scientific and Technical Computing at Lund University, LUNARC, Sweden, partially funded by the Swedish Research Council through grant agreement no. 2018-05973. We acknowledge the National Academic Infrastructure for Supercomputing in Sweden (NAISS), partially funded by the Swedish Research Council through grant agreement no. 2022-06725, for awarding this project access to the LUMI supercomputer, owned by the EuroHPC Joint Undertaking and hosted by CSC (Finland) and the LUMI consortium.

We acknowledge the European Synchrotron Radiation Facility, ESRF, for beamtime MX-2757 (DOI: 10.15151/ESRF-ES-2227803777) and Petra Pernot and Mark Tully for assistance during SAXS data collection.

We thank Elise Pepermans for valuable scientific discussions during the early stages of this work.

## Notes

### Competing Interest Statement

The authors have declared no competing interest.

### Summary of Updates

This version of the manuscript has been revised to update the following: added SASBDB codes.

