## Supporting Information for "The hearing-essential intracellular domain of PCDH15 reveals a new layer of auditory mechanotransduction"

### Supplemental data

**Protein sequence recombinant mouse (CR)CD2:**

**Tag** - HHHHHHHHGGGGSWSHPQFEKGGGSGGGSGGSAWSHPQFEKGGGGSENLYFQG
**CR** - SYRQFKVRQAECTKTARIQSAMPAAKPAAPVPAAPAPPPPPPPPPPGAHLYEELGESAMH
**CD2** - KYEMPQYGSRRRLLPPAGQEEYGEVIGEAEEEYEEEEVEPEKVKKPKVEIREPSEEEVVVTVEKP
PAAEPTYPTWKRARIFPMIFKKVRGLAEKRGIDLEGEEWRRRLDEEDKDYLQLTLDQEEATESTVESEE
ESSDYTEYTETESEFSESETTEESESETPSEEAEESSTPESEESESTESEGEKARKNIVLARRRPVVEEIQE
VKGKREEPPVEEEEEPPLEEEERAEEGEESEAAPMDESTDLEAQDVPEEGSAESVSMERGVESEESES
ELSSSSSTSESLSGGPWGFQVPEYDRRKDEEPKKSPGANSEGYNTAL

#### Supplementary figures

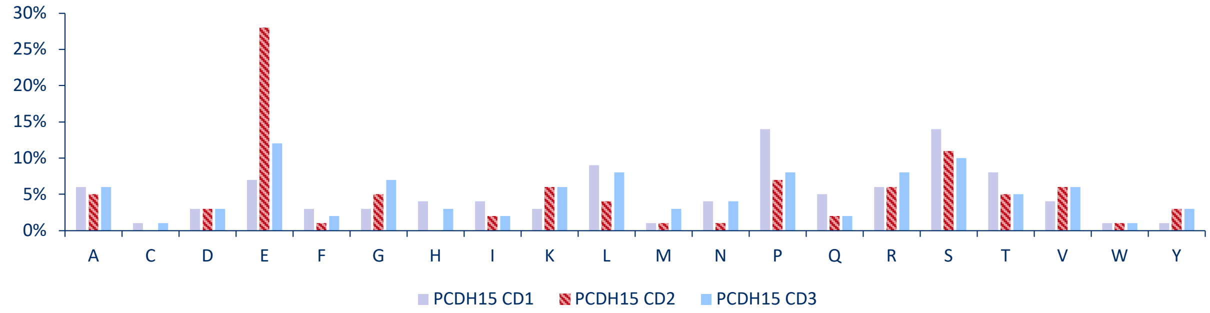


**Supplementary Figure 1 | Amino-acid composition of the PCDH15 cytoplasmic isoforms.** Relative abundance of the 20 amino acids in PCDH15-CD1, PCDH15-CD2 and PCDH15-CD3, calculated from the full cytoplasmic isoform sequences including the common region.

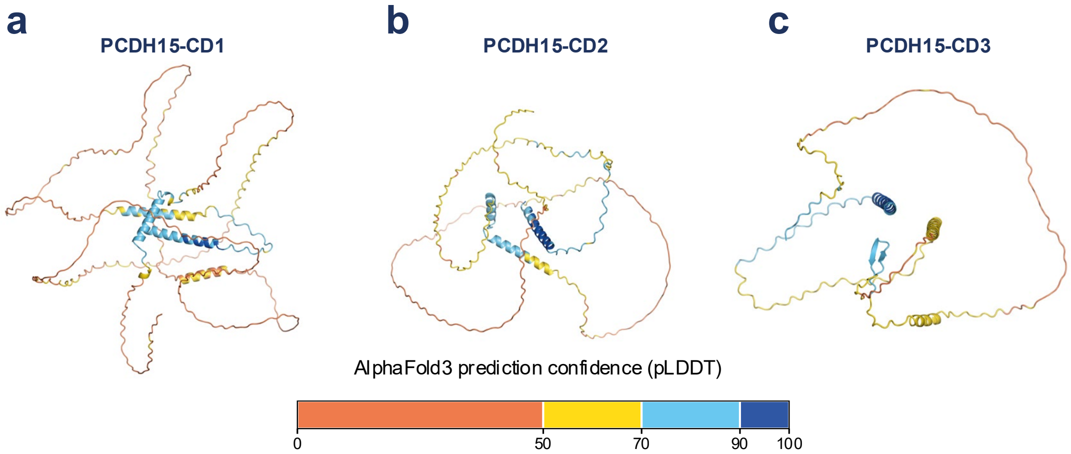


**Supplementary Figure 2 | AlphaFold3 predictions of the PCDH15 cytoplasmic isoforms.** AlphaFold3 predictions of **a,** PCDH15-CD1, **b,** PCDH15-CD2 and **c,** PCDH15-CD3, coloured according to per-residue predicted Local Distance Difference Test (pLDDT) scores. Low pLDDT scores predominate across all three isoforms, with higher-confidence predictions restricted to short regions.

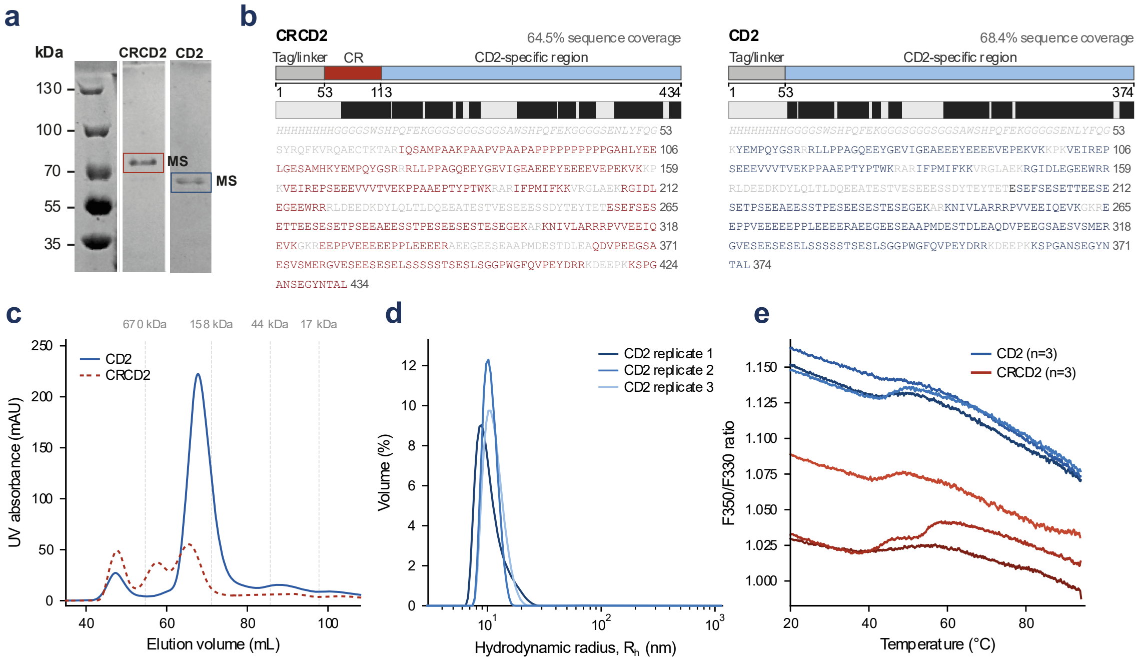


**Supplementary Figure 3 | Biophysical characterisation and validation of recombinant CRCD2 and CD2. (a)** Representative SDS–PAGE of purified CRCD2 and CD2. Boxes indicate the bands excised for identification by LC–MS/MS. **(b)** LC–MS/MS sequence coverage of CRCD2 (left) and CD2 (right). Identified peptides covered 64.5% and 68.4% of the recombinant CRCD2 and CD2 sequences, respectively. Black regions in the coverage tracks indicate residues covered by at least one identified peptide; covered residues are highlighted in red for CRCD2 and blue for CD2 in the corresponding amino-acid sequences. Construct schematics indicate the tag/linker, common region (CR), and CD2-specific region. **(c)** Size-exclusion chromatography (SEC) profiles of CRCD2 and CD2, with the elution positions of globular molecular-mass standards indicated by dashed lines. **(d)** Volume-weighted dynamic light scattering (DLS) size distributions for three CD2 replicates, plotted as hydrodynamic radius ($R_h$). **(e)** Nano-differential scanning fluorimetry (nanoDSF) profiles of CD2 and CRCD2, shown as the F350/F330 fluorescence ratio as a function of temperature (n = 3 replicate measurements per construct). CD2 corresponds to the three upper traces and CRCD2 to the three lower traces.

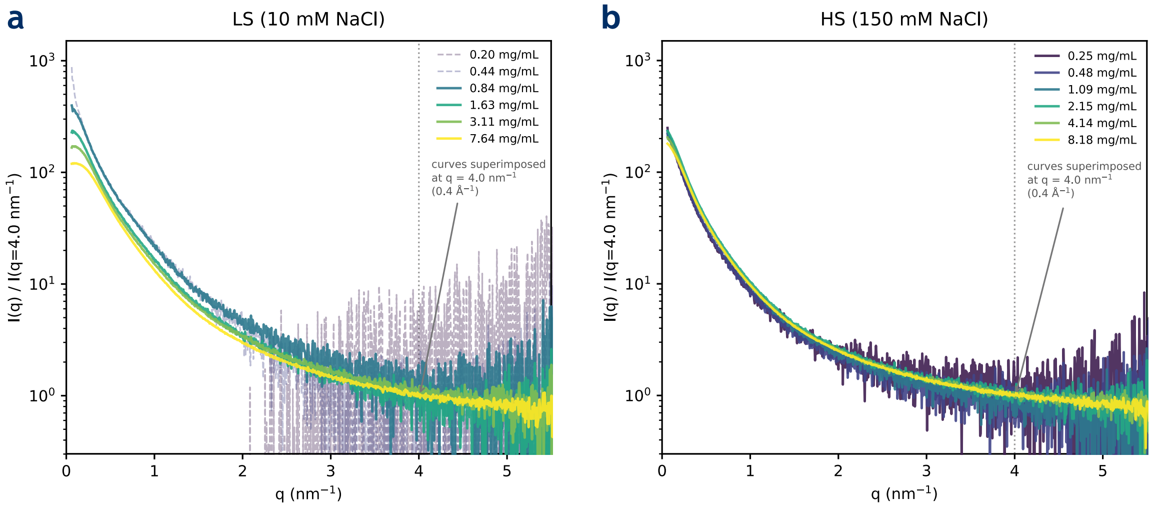


**Supplementary Figure 4 | Concentration dependence of CD2 SAXS profiles.** SAXS profiles collected across six CD2 concentrations in 10 mM NaCl **(a)** and 150 mM NaCl **(b)**. Curves were normalized to the scattering intensity at q = 4.0 $nm^{-1}$ to facilitate comparison of concentration-dependent differences. At 10 mM NaCl, the two lowest-concentration datasets (0.20 and 0.44 mg $ml^{-1}$; dashed lines) showed an additional low-q contribution consistent with trace aggregation and were excluded from concentration extrapolation. The remaining four datasets at 10 mM NaCl and all six datasets at 150 mM NaCl were retained for concentration extrapolation.

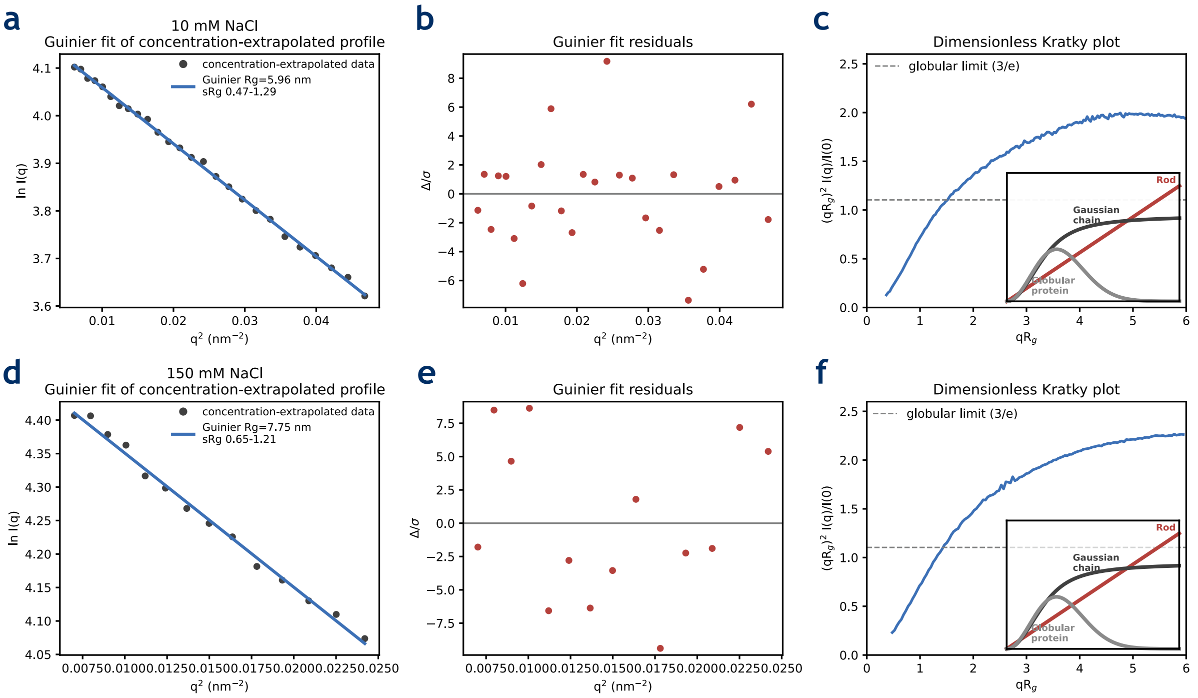


**Supplementary Figure 5 | Guinier and dimensionless Kratky analysis of concentration-extrapolated CD2 SAXS profiles.** Guinier fits (left), corresponding normalized residuals (middle) and dimensionless Kratky plots (right) of the concentration-extrapolated scattering profiles at 10 mM NaCl (top) and 150 mM NaCl (bottom). Guinier analysis yielded Rg values of 5.96 and 7.75 nm, respectively, with the corresponding qRg fitting ranges indicated in the plots. Dashed horizontal lines in the dimensionless Kratky plots indicate the globular reference value of 3/e.

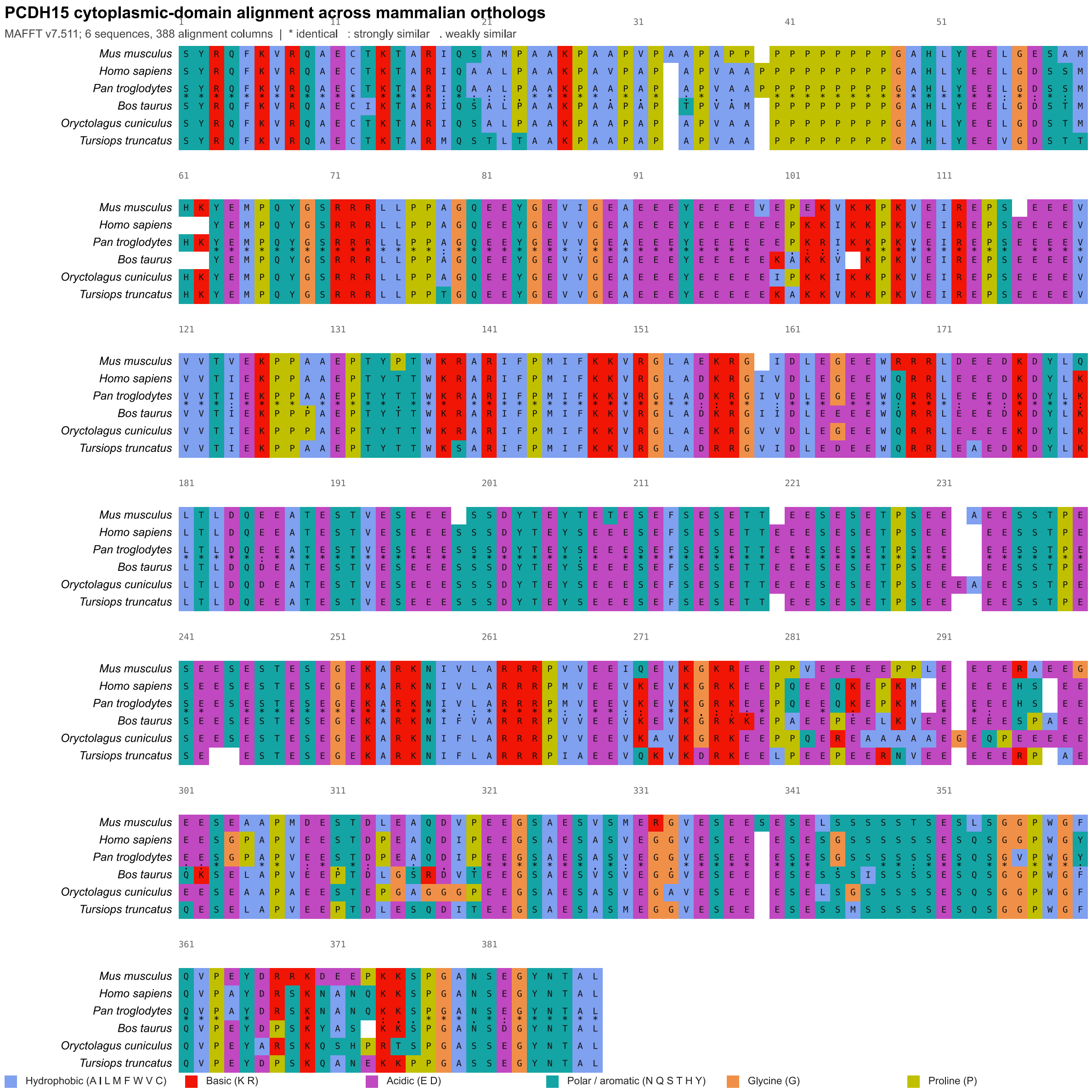


**Supplementary Figure 6 | Sequence conservation of the PCDH15-CD2 cytoplasmic region across mammalian orthologues.** Multiple sequence alignment of the PCDH15-CD2 cytoplasmic region from Mus musculus, Homo sapiens, Pan troglodytes, Bos taurus, Oryctolagus cuniculus and Tursiops truncatus. Residues are coloured according to amino-acid physicochemical class, as indicated in the legend. Residue numbering corresponds to the Mus musculus PCDH15-CD2 cytoplasmic sequence.

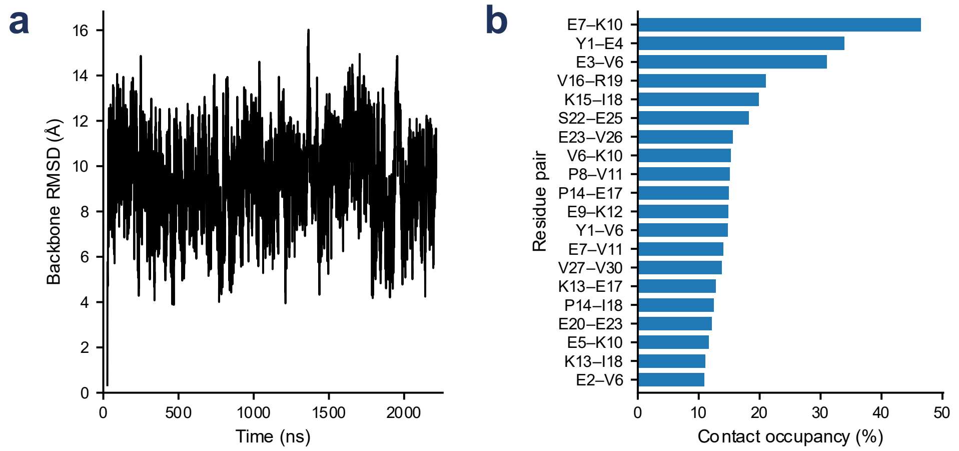


**Supplementary Figure 7 | Conformational variability and transient intramolecular contacts of the simulated CD2 peptide. (a)** Backbone RMSD as a function of time across the combined 3.5-μs analysed trajectory. **(b)** Occupancy of the most frequently observed intramolecular residue–residue contacts across the combined trajectory.

### Supplementary tables

**Supplementary Table 1 | SAXS data collection and structural parameters for the recombinant PCDH15 CD2 intracellular domain.** Parameters were obtained from concentration-extrapolated scattering profiles collected in 10 mM and 150 mM NaCl. Radii of gyration (Rg) were determined independently by Guinier and pair-distance distribution, P(r), analyses; maximum particle dimensions (Dmax) were obtained from the P(r) distributions.

| **Parameter** | **10 mM NaCl** | **150 mM NaCl** |
|---|---|---|
| Beamline | ESRF BM29 | |
| Energy (keV) | 12.5 | |
| Wavelength (Å) | 0.992 | |
| Temperature (°C) | 20 | |
| q-range ($Å^{-1}$) | 0.00776–0.495 | |
| Concentration range (mg $mL^{-1}$) | 0.20–7.64 | 0.25–8.18 |
| Concentrations measured | 6 | 6 |
| Concentrations used for extrapolation | 4 | 6 |
| Guinier *Rg*, c→0 (nm) | 5.96 | 7.75 |
| (*P(r)*) *Rg* (nm) | 6.50 | 8.76 |
| Dmax (nm) | 23.90 | 36.51 |
| MW, theoretical tagged CD2 (kDa) | 42.4 | |
| CorMap comparison (10 vs 150 mM NaCl) | C=764, p=$2.5x10^{-228}$ | |

**Supplementary Table 2 | Predicted short linear motifs within the PCDH15-CD2 intracellular domain.** Candidate motifs were identified using the Eukaryotic Linear Motif (ELM) resource. Conservation was assessed by multiple sequence alignment of six mammalian PCDH15-CD2 orthologues, using mouse PCDH15 as the reference sequence. Sequence conservation was calculated as the percentage of motif positions that were identical across all six mammalian orthologues analysed. Overlapping predictions of the same functional category were merged where appropriate into contiguous motif regions for visualization in Fig. 4 and presentation in this table. Motifs represent computational predictions and do not imply experimentally validated interactions or phosphorylation events.

| Functional category | ELM identifier | Sequence | Residues | Conservation across orthologues |
|---|---|---|---|---|
| SH3-domain ligand (proline-rich) | LIG_SH3_3 | AAKPAAPVPAAPA | 24–36 | 62% invariant (8/13 positions; 6/6 orthologues) |
| Profilin ligand (poly-proline) | LIG_PROFILIN_1 | PPPPPPPPPPG | 37–47 | 82% invariant (9/11 positions; 6/6 orthologues) |
| SH3-domain ligand (proline-rich) | LIG_SH3_3 | PEKVKKP | 100–106 | 29% invariant (2/7 positions; 6/6 orthologues) |
| SH3-domain ligand (proline-rich) | LIG_SH3_3 | VVTVEKP | 119–125 | 86% invariant (6/7 positions; 6/6 orthologues) |
| SH3-domain ligand (proline-rich) | LIG_SH3_3 | AEPTYP | 128–133 | 83% invariant (5/6 positions; 6/6 orthologues) |
| Calcineurin (PP2B) docking motif, PxIxIT-type | DOC_PP2B_PxIxIT_1 | KPKVEIR | 105–111 | 100% invariant (7/7 positions; 6/6 orthologues) |
| MAPK docking motif (generic) | DOC_MAPK_gen_1 | KVKKPKVEI | 102–110 | 67% invariant (6/9 positions; 6/6 orthologues) |
| MAPK docking motif (MEF2A-type) | DOC_MAPK_MEF2A_6 | KVKKPKVEI | 102–110 | 67% invariant (6/9 positions; 6/6 orthologues) |
| MAPK docking motif (generic) | DOC_MAPK_gen_1 | KRARIFPMIF | 136–145 | 90% invariant (9/10 positions; 6/6 orthologues) |
| MAPK docking motif (MEF2A-type) | DOC_MAPK_MEF2A_6 | KDYLQLTL | 173–180 | 88% invariant (7/8 positions; 6/6 orthologues) |
| MAPK docking motif (NFAT4-type) | DOC_MAPK_NFAT4_5 | KDYLQLTLD | 173–181 | 89% invariant (8/9 positions; 6/6 orthologues) |
| MAPK docking motif (generic) | DOC_MAPK_gen_1 | KARKNIVL | 247–254 | 75% invariant (6/8 positions; 6/6 orthologues) |
| TRAF6 ligand motif | LIG_TRAF6_MATH_1 | ETPSEEAE | 221–228 | 88% invariant (7/8 positions; 6/6 orthologues) |
| TRAF6 ligand motif | LIG_TRAF6_MATH_1 | EPPVEEEEEP PLEEEER | 273–289 | 24% invariant (4/17 positions; 6/6 orthologues) |
| PDZ-domain ligand (Class I, C-terminal) | LIG_PDZ_Class_1 | GYNTAL | 376–381 | 100% invariant (6/6 positions; 6/6 orthologues) |
| CK2 phosphorylation site | MOD_CK2_1 | REPSEEE | 111–117 | 100% invariant (7/7 positions; 6/6 orthologues) |
| CK2 phosphorylation site | MOD_CK2_1 | ATESTVESEEE | 185–195 | 100% invariant (11/11 positions; 6/6 orthologues) |
| GSK3 phosphorylation site | MOD_GSK3_1 | ATESTVESEEE SSDYTEYTETE SEFSESETTEES | 185–224 | 95% invariant (38/40 positions; 6/6 orthologues) |

| | | | | |
|---|---|---|---|---|
| | | ESETPS | | |
| Plk1 phosphorylation site | MOD_Plk_1 | TESTVES | 186–192 | 100% invariant (7/7 positions; 6/6 orthologues) |
| CK1 phosphorylation site | MOD_CK1_1 | SDYTEYT | 197–203 | 86% invariant (6/7 positions; 6/6 orthologues) |
| CK2 phosphorylation site | MOD_CK2_1 | TEYTETESEFSE SETTEESESETPSE | 200–225 | 92% invariant (24/26 positions; 6/6 orthologues) |
| CK1 phosphorylation site | MOD_CK1_1 | SEFSESETTEES | 207–218 | 100% invariant (12/12 positions; 6/6 orthologues) |
| Plk2/3 phosphorylation site | MOD_Plk_2-3 | ETPSEEA | 221–227 | 86% invariant (6/7 positions; 6/6 orthologues) |
| CK2 phosphorylation site | MOD_CK2_1 | EESSTPESEEES TESEGE | 228–246 | 89% invariant (17/19 positions; 6/6 orthologues) |
| CK1 phosphorylation site | MOD_CK1_1 | SEESESTESEGE | 235–246 | 83% invariant (10/12 positions; 6/6 orthologues) |
| CK2 phosphorylation site | MOD_CK2_1 | DESTDLE | 302–308 | 29% invariant (2/7 positions; 6/6 orthologues) |
| Plk2/3 phosphorylation site | MOD_Plk_2-3 | DESTDLE | 302–308 | 29% invariant (2/7 positions; 6/6 orthologues) |
| CK1 phosphorylation site | MOD_CK1_1 | SAESVSM | 317–323 | 71% invariant (5/7 positions; 6/6 orthologues) |
| Plk2/3 phosphorylation site | MOD_Plk_2-3 | ESVSMER | 319–325 | 57% invariant (4/7 positions; 6/6 orthologues) |
| GSK3 phosphorylation site | MOD_GSK3_1 | ESEESESELSS SSSTSESLS | 328–347 | 65% invariant (13/20 positions; 6/6 orthologues) |
| CK1 phosphorylation site | MOD_CK1_1 | SEESESELSSS SSTSESL | 329–346 | 61% invariant (11/18 positions; 6/6 orthologues) |
| CK2 phosphorylation site | MOD_CK2_1 | SEESESE | 329–335 | 86% invariant (6/7 positions; 6/6 orthologues) |
| CK2 phosphorylation site | MOD_CK2_1 | SSSSTSE | 338–344 | 57% invariant (4/7 positions; 6/6 orthologues) |
| Phospho-dependent docking motif (Pin1 WW-domain) | DOC_WW_Pin1_4 | ESETPS | 219–224 | 100% invariant (6/6 positions; 6/6 orthologues) |
| Phospho-dependent docking motif (Pin1 WW-domain) | DOC_WW_Pin1_4 | ESSTPE | 229–234 | 100% invariant (6/6 positions; 6/6 orthologues) |
| Phospho-dependent docking motif (Pin1 WW-domain) | DOC_WW_Pin1_4 | PKKSPG | 366–371 | 33% invariant (2/6 positions; 6/6 orthologues) |
